# Morphological map of under- and over-expression of genes in human cells

**DOI:** 10.1101/2024.12.02.624527

**Authors:** Srinivas Niranj Chandrasekaran, Eric Alix, John Arevalo, Adriana Borowa, Patrick J. Byrne, William G. Charles, Zitong S. Chen, Beth A. Cimini, Boxiong Deng, John G. Doench, Jessica D. Ewald, Briana Fritchman, Colin J. Fuller, Jedidiah Gaetz, Amy Goodale, Marzieh Haghighi, Yu Han, Zahra Hanifehlou, Holger Hennig, Desiree Hernandez, Christina B. Jacob, Tim James, Tomasz Jetka, Alexandr A. Kalinin, Ben Komalo, Maria Kost-Alimova, Tomasz Krawiec, Brittany A. Marion, Glynn Martin, Nicola Jane McCarthy, Lisa Miller, Arne Monsees, Nikita Moshkov, Alán F. Muñoz, Arnaud Ogier, Magdalena Otrocka, Krzysztof Rataj, David E. Root, Francesco Rubbo, Simon Scrace, Douglas W. Selinger, Rebecca A. Senft, Peter Sommer, Amandine Thibaudeau, Sarah Trisorus, Rahul Valiya Veettil, William J. Van Trump, Sui Wang, Michał Warchoł, Erin Weisbart, Amélie Weiss, Michael Wiest, Agata Zaremba, Andrei Zinovyev, Shantanu Singh, Anne E. Carpenter

## Abstract

Cell Painting images offer valuable insights into a cell’s state and enable many biological applications, but publicly available arrayed datasets only include hundreds of genes perturbed. The JUMP (Joint Undertaking in Morphological Profiling) Cell Painting Consortium perturbed roughly 75% of the protein-coding genome in human U-2 OS cells, generating a rich resource of single-cell images and extracted features. These profiles capture the phenotypic impacts of perturbing 15,243 human genes, including overexpressing 12,609 genes (using open reading frames, ORFs) and knocking out 7,975 genes (using CRISPR-Cas9). We mitigated technical artifacts by rigorously evaluating data processing options and validated the dataset’s robustness and biological relevance. Analysis of phenotypic profiles revealed novel gene clusters and functional relationships, including those associated with mitochondrial function, cancer, and neural processes. The JUMP Cell Painting genetic dataset is a valuable resource for exploring gene relationships and uncovering novel functions.

## Introduction

A major biological challenge is to identify the functions of all human proteins and to understand the impact of disease on those functions. Historically, human protein functions are determined painstakingly, often through hypothesis-driven and heavily customized genetic or proteomic experiments. The pace of discovery increased with the rise of genome-scale perturbation screens that simultaneously assess all human proteins for a given function. First in arrayed format and later in a more economical pooled format, these experiments capture readouts (reflecting phenotypes of interest) from reporter assays, molecular -omics, or microscopy. They can identify all genetic perturbations that impact a single or a few biological processes of interest, within limits of technical noise ^1–4^.

As technologies increased the *scale* of functional genomics experiments, they also increased the *breadth* of readouts measured from a sample. Proteomic, metabolomic, and transcriptomic techniques allow measuring hundreds to thousands of individual molecular readouts. Profiling technologies enable rapid discovery of new protein functions in two ways. First, researchers can identify, in a single experiment, all genes whose perturbation impacts each of the hundreds of individual direct readouts in the assay. Second, an entirely new mode of discovery arose: using the complex patterns among the hundreds of readouts as a “profile” or signature to assign novel functions to genes based on “guilt-by-similarity”.

Images of cells captured by fluorescence microscopy have emerged as a surprisingly powerful profiling methodology alongside molecular profiling. Image analysis extracts thousands of morphological features ^5^ and captures biological information on par with available high-throughput proteomic or transcriptomic methods ^6,7^, at single-cell resolution ^8–10^. Using images to generate profiles has proven useful for identifying gene functions, determining the mechanism of compounds, and identifying novel chemical regulators of genes, among many other applications in drug discovery and fundamental biological research ^11^.

Accordingly, information-rich imaging datasets have been publicly shared ^12^ that capture morphological profiles of thousands of genetic perturbations. An early Mitocheck consortium screen decreased the expression of 21,000 human genes by RNA interference, capturing the impact on chromosome morphology in time-lapse ^13^. Multiple studies using deletion strains in yeast have captured morphological impact ^14,15^. Morphological profiles of actin, DNA, and α-tubulin staining were captured for 6,840 Drosophila genes perturbed by RNA interference, plus pairwise combinations of those genes with a 100-gene subset ^16^.

Cell Painting has become fairly standard as an image-based profiling readout. Thus far, the largest arrayed genetic perturbation dataset using the assay is from the techbio company Recursion, using six CRISPR-Cas9 guides against 736 genes in human umbilical vein endothelial cells (HUVEC) (with an additional 16,327 genes anonymized) ^17^. New methods allow pooling thousands of genetic reagents in human cells while still offering image-based readouts; we helped execute three Cell Painting screens using CRISPR-Cas9 guides to knock down each of >20,000 human genes, though an average of only 460 cells’ data are available per gene (<100 per guide), and no data from overexpression perturbations is available ^18^.

Here, we aimed to create a public, large-scale dataset of Cell Painting image-based profiles from increasing and decreasing gene expression, in an arrayed format that allows thousands of cells per replicate. Over- and under-expression of proteins can both have advantages for impacting cell systems and suggesting biological functions: unlike knockdown/knockout technologies, overexpression can produce phenotypes even in (perhaps, especially in) a cell type where the gene isn’t expressed or when there are multiple isoforms or highly redundant protein family members. Overexpression does not suffer from off-target effects and chromosome arm effects that can impact CRISPR-Cas9 systems ^19^ but is prone to artifactual effects — though some, such as dominant negative effects, may still affect a biological function specifically enough to be informative ^20^. This dataset is publicly shared as the genetic subset of the 136,000 chemical and genetic perturbations tested by our Joint Undertaking for Morphological Profiling (JUMP) Cell Painting Consortium ^21^.

## Results

### Gene overexpression and CRISPR-Cas9 knockout yield Cell Painting image-based phenotypes

Using the Cell Painting assay v3 ^22^, we tested 15,142 overexpression (“ORF”) reagents encompassing 12,609 unique human genes, including controls from the Broad Institute lentiviral ORF library ^23^. We also tested 7,977 pools of CRISPR guides against 7,975 unique genes and controls from the Human Edit-R^TM^ synthetic crRNA - Druggable Genome library from Revvity Discovery Limited (formerly known as Horizon Discovery). In total, we tested 15,243 genes by overexpression, knockout, or both. Lentiviruses containing each ORF, or pools of four CRISPR guides against a given gene, were arrayed into 384-well plates of U-2 OS human osteosarcoma cells. The cells were incubated for 48 hours to allow infection (or transfection) and expression (or transient knockout) and then fixed and stained for the Cell Painting assay, with six stains labeling eight cellular components or organelles and imaged in five fluorescent channels (Figure 1 a). The design of the experiment is shown in Figure 1 b and described in detail in the Methods section. We extracted 7,648 and 4,761 image-based features from the ORF and CRISPR datasets, respectively (https://github.com/jump-cellpainting/2024_Chandrasekaran_Morphmap/tree/main/11.list-all-features/output). The features include metrics of size, shape, intensity, texture, and correlation, measured from each of the channels and in the nucleus, cytoplasm, and cell compartments. Quality control, feature selection, and other pre-processing steps (see Methods) yielded final profiles with 722 and 259 features for ORF and CRISPR profiles, respectively. Notably, the sphering and harmony steps used in pre-processing yield final features that cannot be readily mapped to the original feature categories, making interpretability challenging.

**Figure 1:**
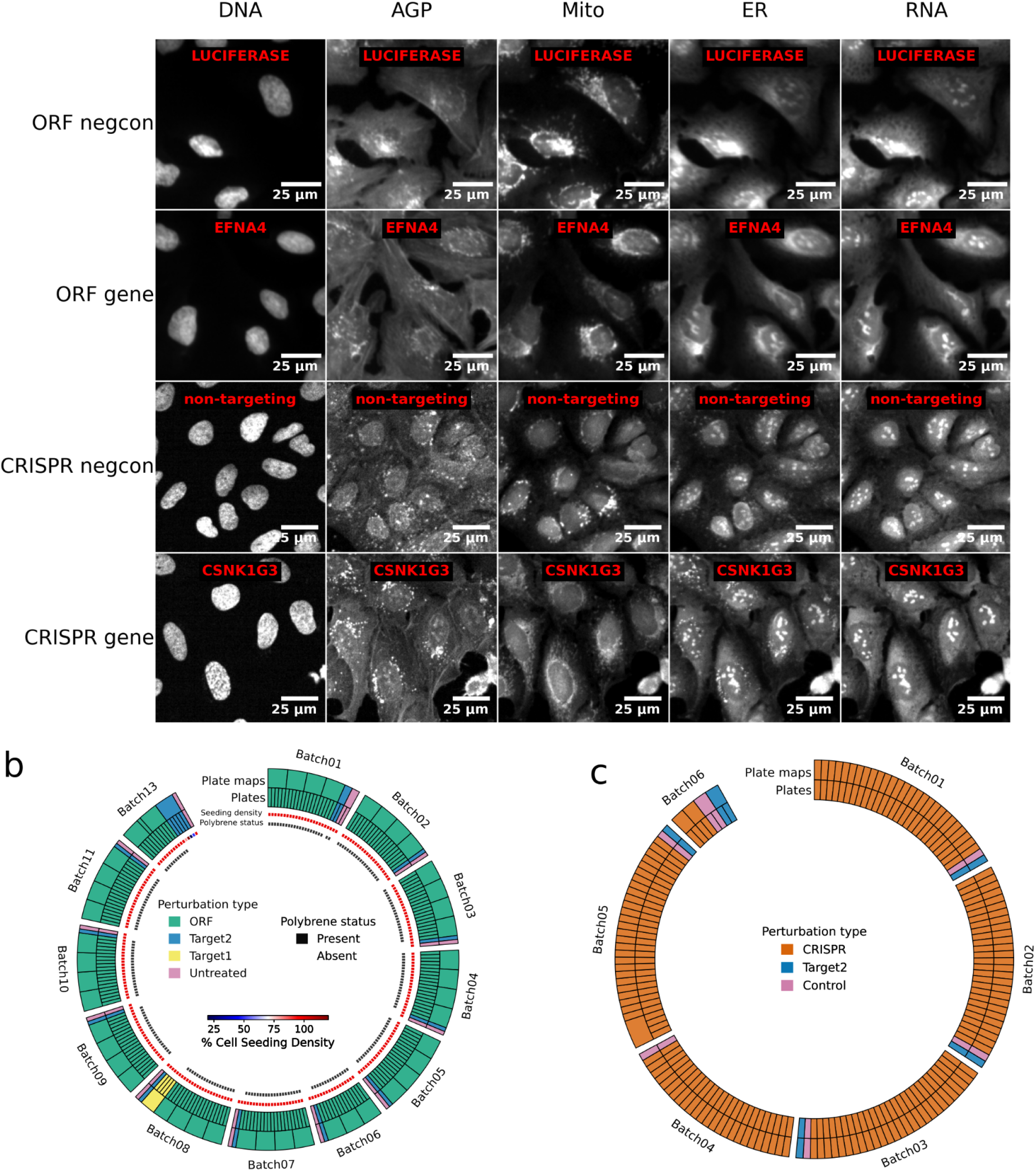
JUMP Cell Painting dataset overview. a) Example images of overexpressed genes (ORFs) and knocked-out genes (CRISPR) that are most dissimilar from negative controls (negcon) are shown. Negative control images are selected randomly from the ORF and CRISPR dataset. Channels are DNA: nucleus (Hoechst); AGP: actin cytoskeleton (phalloidin) plus the Golgi and plasma membrane (wheat germ agglutinin); Mito: mitochondria (MitoTracker); ER: endoplasmic reticulum (concanavalin A); RNA: nucleoli and cytoplasmic RNA (SYTO 14). b) and c) Experimental design schematic for dataset generation. The number of plates, plate maps and batch in the ORF and CRISPR datasets are shown. Detailed information regarding perturbation types per plate and other experimental conditions (e.g., cell seeding density) can be found in the Methods section.

Overall, of the 15,243 genes tested (Figure 2 a), 68% (10,352) yielded a detectable phenotype (“phenotypic activity”), that is, an image-based profile with a signal distinct from negative controls by ORF, CRISPR, or both ^24^. This includes 7,031 genes (56% of tested genes) by overexpression and 5,546 genes (70% of tested genes) by CRISPR-Cas9 knockout (Figure 2 b and c Supplementary Table 1). Clustering revealed many groups with known relationships (Supplementary Figure 1).

**Figure 2:**
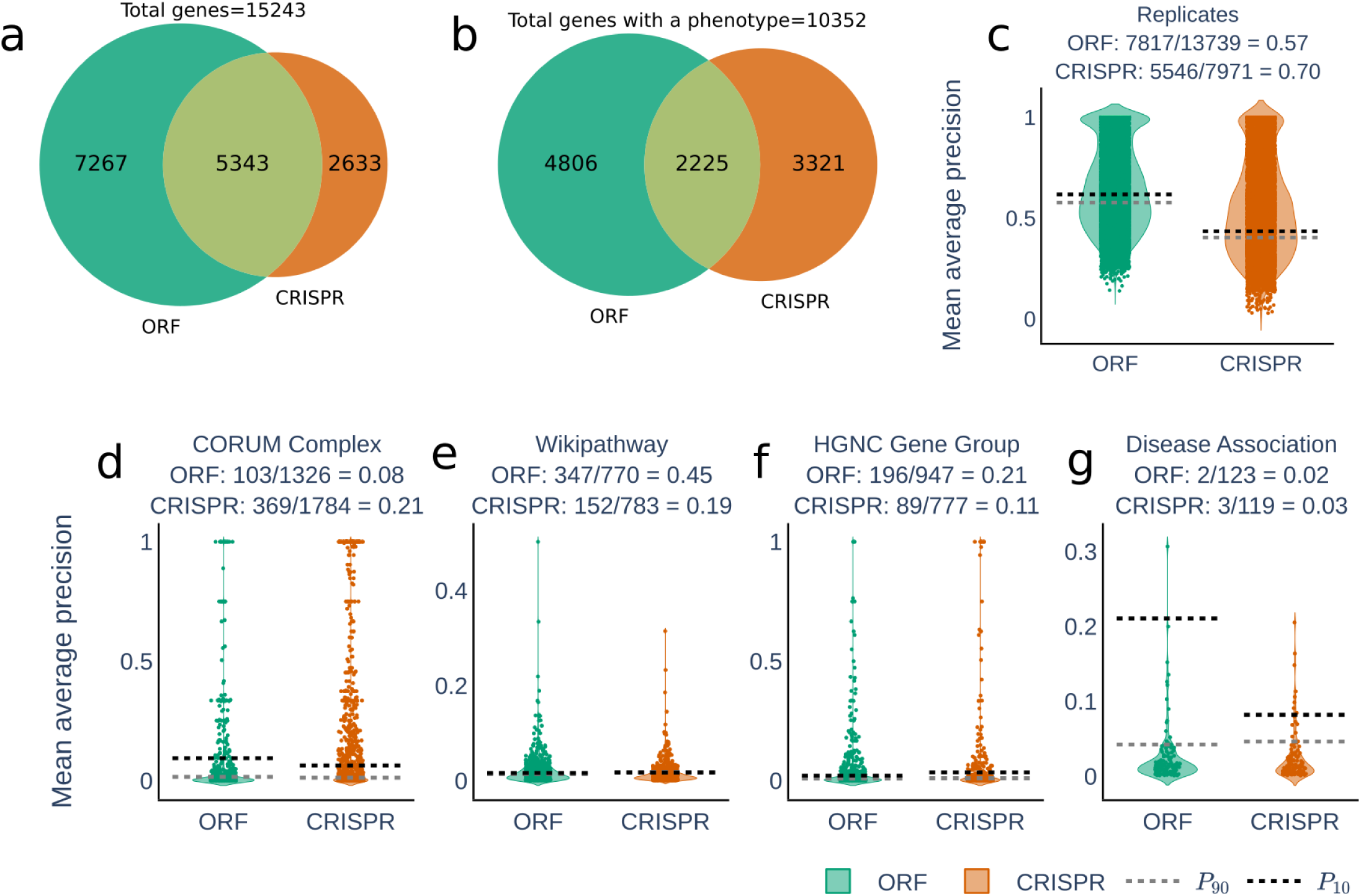
Phenotypic activity for genes and phenotypic consistency for gene groups. a) Number of genes overexpressed, knocked-out, or both. The Broad Institute lentiviral ORF library is not genome-wide because genes above 3,500 nucleotides do not package well into lentivirus, and some genes were excluded due to quality control reasons. b) 68% of the tested genes yielded a phenotype (replicates are significantly different from the negative control); 56% of genes had a phenotype by ORF and 70% by CRISPR. c) To determine whether a gene had a detectable phenotype (“phenotypic activity”), we computed mean average precision (mAP) ^24^ to assess how well a given replicate of a gene retrieves other replicates of that gene rather than a negative control. The remaining plots (d-g) display phenotypic consistency, as measured by the mAP for retrieval of genes sharing specific labels/categories rather than replicates or genes with different labels/categories, based on human knowledge (CORUM Complex — also shown in Supplementary Figure 2, WikiPathways, HGNC Gene Group, or Disease Association — details in Methods). Green: ORF dataset; Orange: CRISPR dataset. The fraction of genes/gene groups with significant mAP values is shown above each panel. The gray and black dashed lines, respectively, are the 90th percentile of the genes/gene groups with a non-significant mAP value and the 10th percentile of genes/gene groups with a significant mAP value (the thresholds vary because there are varying numbers of replicates per gene or genes per group).

### Genes likely to have Cell Painting phenotypes share certain characteristics

We performed Fisher’s exact test to discover whether particular types of genes were more likely to have phenotypic activity in Cell Painting (Figure 3 a). As expected, essential genes are more likely than random genes to have a phenotype when knocked out and less likely when overexpressed; essential genes as a class are already well-expressed, so overexpression may not dramatically impact the total amount of protein.

**Figure 3:**
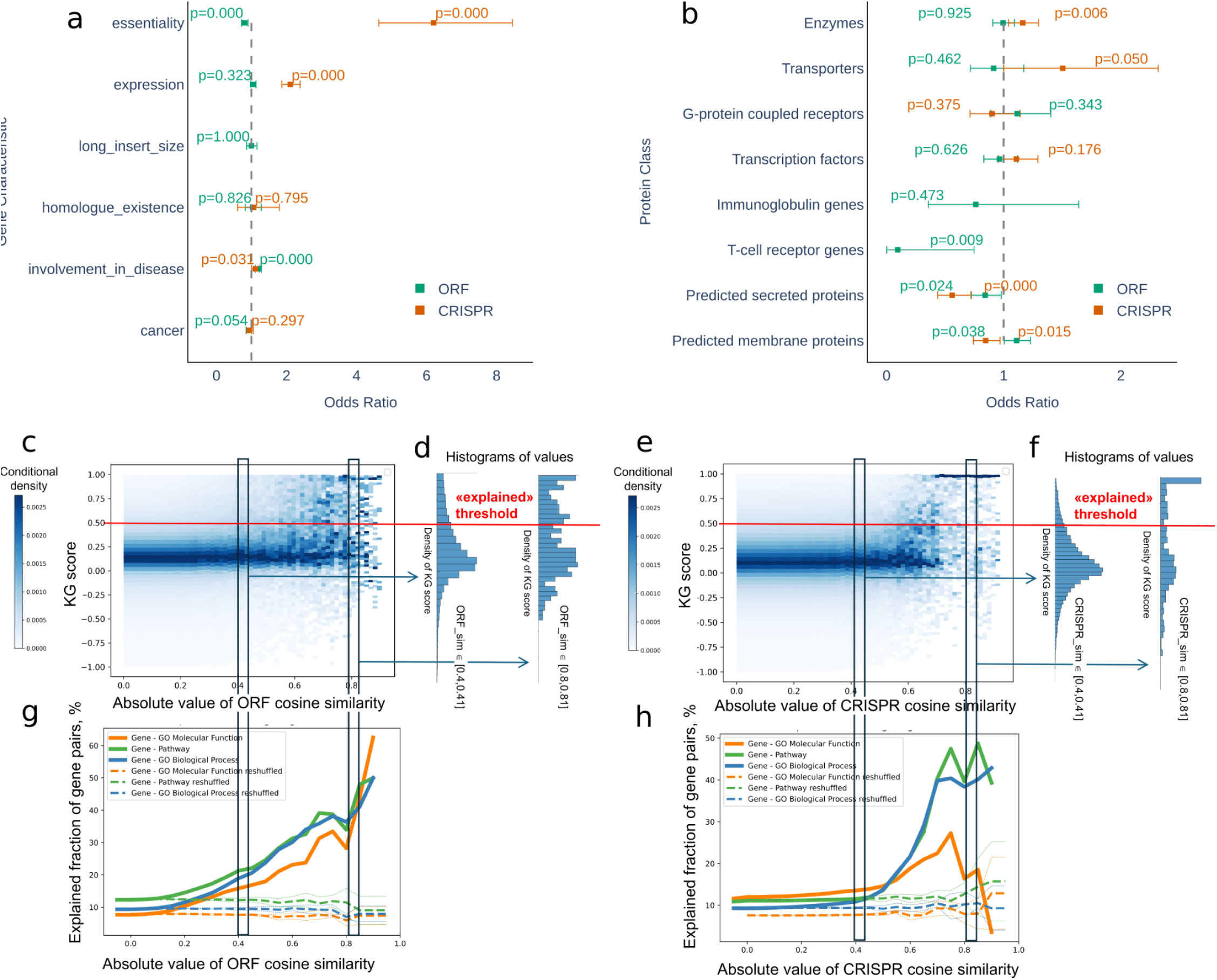
Characteristics of genes with phenotypes and knowledge graph validation of the dataset. Forest plots illustrating associations between gene characteristics/protein classes and observed phenotypes. Green represents ORF profiles, and orange represents CRISPR profiles. Odds ratios (ORs), confidence intervals, and significance levels (calculated using Fisher’s exact test) are displayed. Multi-hypothesis correction resulted in adjusted significance thresholds (α = 0.0083 for a and α = 0.0062 for b). Roughly half of gene-gene connections in Cell Painting are validated by existing knowledge from DRKG. c-f) Comparison of morphological similarity between gene pairs in JUMP Cell Painting with evidence from the Drug Repurposing Knowledge Graph (DRKG). For ORF profiles (c) and CRISPR profiles (f), the two-dimensional (2D) histogram compares the morphological similarity between gene pairs (x-axis, absolute value of cosine similarity between image-based profiles) with the extent of existing knowledge about that connection (y-axis, KG score; we define a threshold of 0.5 above which a link is considered “explained” through the KG’s GNN prediction). Data points trend towards the upper right, indicating gene-gene pairs with strong image-based similarity are more likely to have existing evidence in the knowledge graph than morphologically unrelated pairs. The large number of gene pairs is better depicted as a 2D histogram than a scatter plot. For ORF profiles (d) and CRISPR profiles (g), pairs of one-dimensional histograms are shown as examples, which are cuts through the 2D histogram in (a/d); the left of each pair of histograms includes gene pairs in the range [0.4,0.41] (moderate morphological similarity), and the right histogram includes gene pairs in the range [0.8,0.81] (strong morphological similarity). The gene pairs with stronger morphological similarity are more likely to have stronger KG scores. For ORF profiles (g) and CRISPR profiles (h), about 40-50% of the gene pairs with strong morphological similarity (absolute value >0.8) can be explained from the knowledge graph, i.e., the gene-gene links are predicted with the GNN. This view of the data shows three variations of the knowledge graph, trained on molecular functions (gene_mf), pathways (gene_pathways), and biological processes (gene_bp). A random shuffling of gene labels shows that around 10% of gene pairs have connections if selected randomly (dashed lines show standard confidence intervals corresponding to one standard deviation for multiple shuffling of gene labels). Gene pairs with morphological similarities above ∼0.4 (absolute value) show more DRKG evidence for connection than random pairs (except for the GNN trained with molecular function GO terms for CRISPR data).

Likewise, knocking out highly expressed genes is more likely to yield phenotypic impact than overexpressing such genes. Over-expressing disease-associated genes were more likely than non-disease-related genes to produce phenotypic changes. This has implications for drug discovery, as small molecules that reverse these phenotypic changes might be pursued as disease therapeutics.

We also tested whether particular classes of genes, as defined by The Human Protein Atlas ^25^, were more likely to yield Cell Painting phenotypes (Figure 3 b). For gene knockouts, enzymes were more likely than random genes to have phenotypic activity, while predicted secreted and membrane proteins were less likely.

For overexpression, predicted secreted proteins and T-cell receptors were less likely to have phenotypic activity, while predicted membrane proteins were more likely.

### Cell Painting gene-gene connections are validated by existing knowledge

To assess the value of image-based profiles in representing cell states, we investigated whether genes within the same annotated group were more similar than genes from different groups (Figure 2 d-g). In both the ORF and CRISPR datasets, genes cluster based on their associated CORUM protein complex, WikiPathway, and gene group. Notably, several clusters show phenotypic consistency in both ORF and CRISPR datasets (Supplementary Table 2). Genes do not cluster by disease association, possibly because diverse mechanisms can underpin a single disease.

Next, we compared gene-gene morphological connections to the Drug Repurposing Knowledge Graph (DRKG), which integrates multiple sources of existing connections among genes^26^ (https://github.com/gnn4dr/DRKG). DRKG includes protein-protein interactions, gene co-expression, and other gene-gene links derived from text mining, gene ontology, gene pathway, gene-disease, and other annotations, but it was *not* built using the JUMP Cell Painting (nor indeed any image-based information). We analyzed the closeness of genes in the knowledge graph with a graph neural network (GNN) approach, where links between genes and functions in DRKG are predicted by the GNN and summarized in a GNN-based knowledge graph score (see Methods).

Gene-gene pairs with strong morphological similarity (or strong dissimilarity) were much more likely to have strong existing evidence of a relationship in the knowledge graph (Figure 3 c-h). Roughly half of such gene-gene pairs were supported by existing scientific evidence, as captured by the knowledge graph, and roughly half were novel connections (whereas only 10% of randomly shuffled gene pairs had supporting evidence). The JUMP dataset therefore contains reliable data and also the opportunity for discovery.

### Screening for a single-cell phenotype using machine learning

Cell Painting images contain multiple channels and can reveal hundreds of morphological phenotypes of interest to biologists. To demonstrate the potential to use this data to carry out specific screens, we trained a single-cell phenotypic machine-learning model to recognize rounded cells using the PhenoSorter feature in Spring Engine. Using a point-and-click interface, we provided the supervised learning tool with examples of 53 rounded cells and another 179 flat cells. The resulting classification model trained on an 80/20 training/test split dataset yielded 100% recall and 100% precision in correctly calling rounded cells (Figure 4 a). Applying the model to the entire dataset as a virtual screen, we identified 187 genes in the ORF dataset with excess rounded cells, 63 of which had WikiPathways annotations. All the WikiPathways annotation groups significantly enriched in the rounded-cell gene set were related to cell death or stress (Figure 4 d).

**Figure 4:**
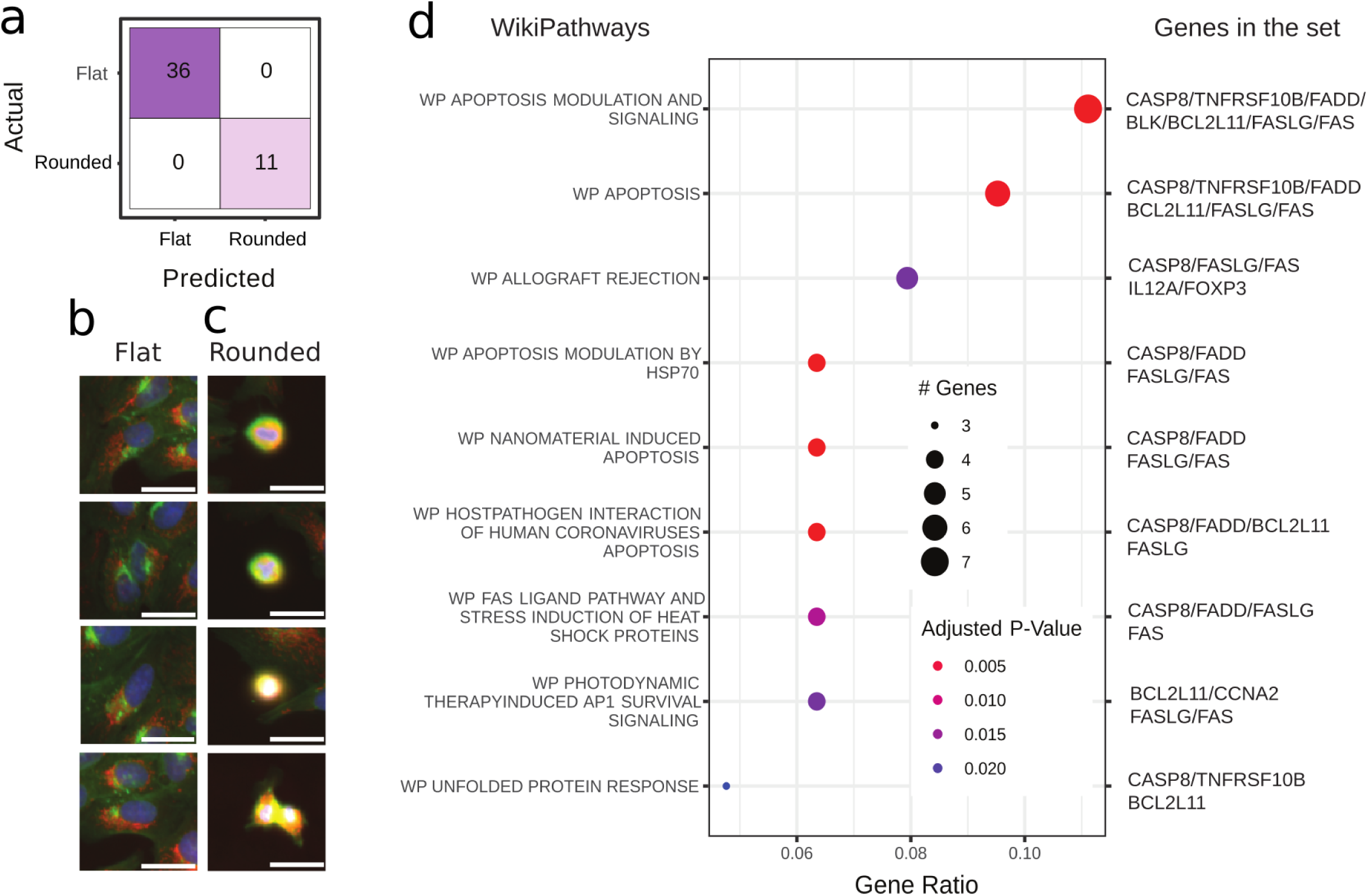
Virtual screen for genes yielding rounded cells. a) Confusion matrix for the single-cell classification model, showing the number of cells in the held-out test set for each combination of Actual vs. Predicted classes. b) Example ‘Flat’ cell image crops from the training set. Scale bars are 10 µm. c) Example ‘Rounded’ cells. d) Dot plot showing significantly enriched gene annotations among genes with a high proportion of ‘Rounded’ cells. Five gene sets were related to apoptosis, and the remaining four pertain to cell stress (individual genes in each set are listed on the right). The dot size reflects the number of hit genes in the indicated gene set. The color indicates the adjusted p-value.

### Identifying gene-gene relationships using image-based profiles

#### Relationships affirmed by literature: Dynein family, FOXO3/TGFb, and SLC39A1/ZBTB16

Given the promising quantitative result that most relationships in the JUMP genetic dataset are supported by literature, we examined some well-known proteins, starting with some of the strongest gene clusters. In the CRISPR dataset, FOXO3 and TGFB1 are negatively correlated (Figure 5 b). This interaction is well known, though the directionality of interaction is tissue-dependent ^27,28^. SLC39A1 and ZBTB16 also strongly negatively correlate in both their ORF and CRISPR profiles (Figure 5 c, d); though lacking direct connection in the literature, the pair shares Gene Ontology annotations, including *Appendage development*, *Embryonic morphogenesis*, and *Skeletal system development*.

**Figure 5:**
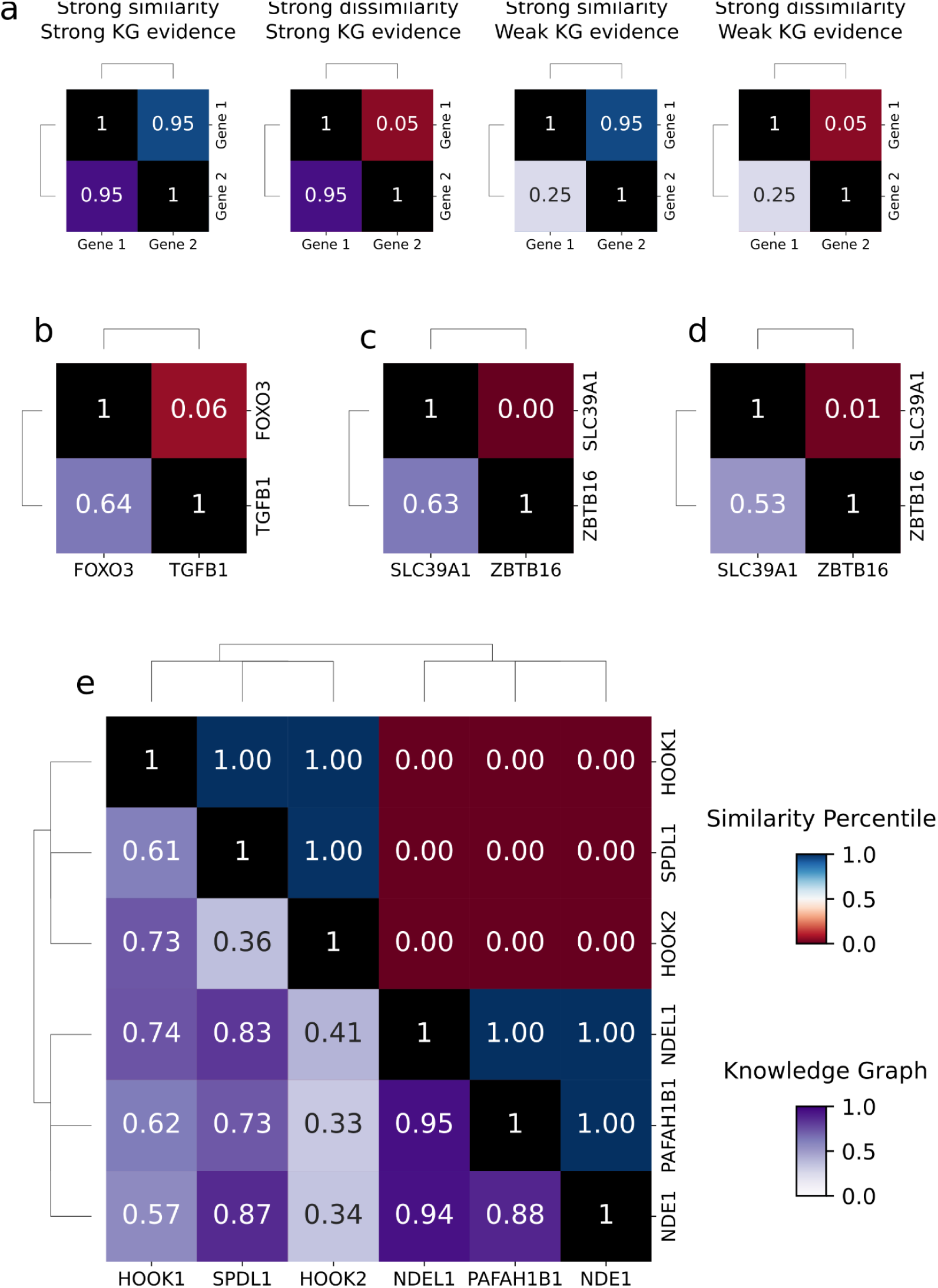
Validation of known gene-gene relationships. a) Schematic representation of a clustergram integrating gene similarity with knowledge graph evidence. The upper triangular matrix displays the percentile of cosine similarity between gene profiles (0 = negative correlation, 1 = positive correlation). The lower triangular matrix shows the knowledge graph score (see Methods), with scores < 0.5 indicating novel associations not predicted by the GNN on DRKG. b-e) Known gene relationships across different perturbation modalities (ORF and CRISPR). b) FOXO3-TGFB1 (CRISPR). c) SLC39A1-ZBTB16 (ORF). d) SLC39A1-ZBTB16 (CRISPR). e) Dynein family cluster (ORF).

In the overexpression (ORF) data, we noticed a strong cluster of PAFAH1B1 (also known as Lis1) with NDE1 (NudE) and NDEL1 (Nudel). HOOK1, HOOK2, and SPDL1 had a strong anti-correlation with those three genes (Figure 5 e). Existing evidence links all six proteins to intracellular dynein transport ^29^; for example, the Nudel complex links PAFAH1B1 to dynein ^30^ and HOOK2 supports the recruitment of dynein by the Nudel complex at the nuclear envelope ^31,32^. As seen in our past work ^33^: both similar and dissimilar morphological profiles can usefully point to functional relationships. “Unexpected” directionality can be due to, for example, biological mechanisms being non-linear, overexpression causing a dominant-negative effect (for example, by disrupting a functioning complex) or feedback loop/compensatory impact on the gene’s function.

We then sought to identify novel discoveries. Again using the DRKG, we prioritized gene-gene connections with strong morphological similarity/dissimilarity but with low existing knowledge per the knowledge graph (using a cutoff of 0.5 for the knowledge graph score, Figure 3). To distinguish whether these are false positives, reflecting technical noise, or true novel relationships between genes, we designed follow-up experiments and were able to make novel discoveries for several of them.

#### MYT1 transcriptionally represses RNF41

One of the top 15 strongest anti-correlating pairs in the CRISPR dataset was MYT1 and RNF41 (Figure 6 a). MYT1 is a known transcriptional repressor ^34^, so we hypothesized it might directly repress transcription of RNF41 — an obvious mechanism that could yield opposite Cell Painting phenotypes. Consistent with this, we performed CUT&RUN analysis and found that MYT1 binds the promoter of RNF41 from adult mouse retinas (Figure 6 f).

**Figure 6:**
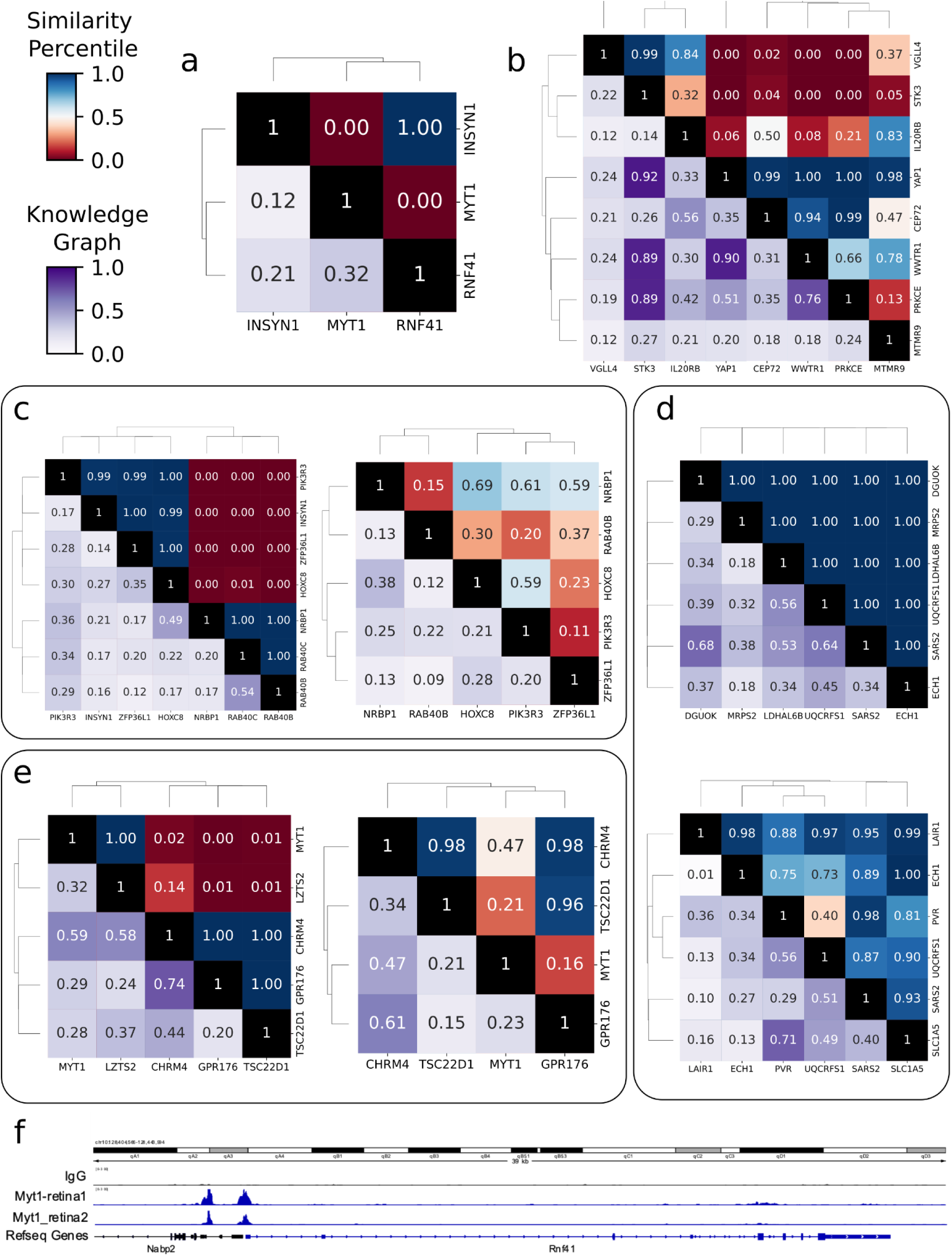
Novel Gene Clusters with Strong Morphological Similarity in JUMP Cell Painting. a - e) Clustergrams highlight novel gene clusters with strong morphological relationships, but with little existing evidence linking the genes. (a) MYT1-RNF41 (ORF). b) A YAP1-associated cluster (ORF). c) Clusters of genes involved in cancer proliferation plus INSYN1 (ORF and CRISPR). d) Clusters of genes implicated in mitochondrial function and cancer (ORF and CRISPR, respectively). e) Clusters of genes involved in neural function (ORF and CRISPR, respectively). Note: RNF41 (from Fig. 6a) is a near neighbor to MYT1 but only appears in a slightly larger version of this cluster. f) Cut & Run demonstrating Myt1 binding at the promoter region of Rnf41 in the mouse retina.

#### Hippo pathway members

We previously noted strong morphological similarities between overexpression of YAP1 and its paralog WWTR1 (TAZ) in an experiment with 220 genes ^33^; we saw the same connection here in the much larger gene set, now with other known Hippo pathway members, STK3 (MST2), VGLL4, and PRKCE ^35,36^ (Figure 6 b).

Using the Plex web application to explore existing datasets (Methods) we found that 10 to 12 of the 50 genes found to be most-similar to YAP1 in the ORF data are co-regulated in various perturbations/conditions, and these genes mostly relate to actin binding/regulation and neuronal development ^37–42^. Additionally, 18 of the YAP1-similar genes exhibit significant expression changes upon YAP1 knockout, further strengthening evidence for a valid biological relationship ^43^. Notably, CEP72, IL20RB, and MTMR9 showed strong phenotypic correlation with YAP1 despite having little knowledge graph evidence for engagement with the Hippo pathway. These genes merit investigation for involvement in the Hippo pathway and related disorders.

#### INSYN1, a potential novel regulator of cell migration and proliferation in cancer

In both the ORF and CRISPR data, many genes linked to the regulation of cell migration and proliferation in cancer cluster together, including the transcription factors HOXC8 and ZFP36L1, the kinase-related genes PIK3R3 and NRBP1, and the GTP-related genes RAB40B and RAB40C (Figure 5 c). INSYN1’s only known function is coordinating the postsynaptic inhibition complex ^44^, but it is also present in this ORF cluster (it was not tested by CRISPR). We found multiple sources of weak evidence for a link between INSYN1 and cancer. For example, the Human Protein Atlas re-analyzed the TCGA dataset and classified INSYN1 as a prognostic marker gene for renal cancer and as having enhanced expression in glioma cancer tissues (https://www.proteinatlas.org/ENSG00000205363-INSYN1/pathology). The Genetic Associations Database contains associations between uncharacterized proteins and cancer outcomes and flagged INSYN1 (under its synonym C15orf59) as one of 62 novel OncoORFs ^45^. Expression of the antisense lncRNA of INSYN1 is associated with low-grade glioma prognosis ^46^ and with the overexpression of vimentin, a marker of advanced metastatic cancer ^47^. Collectively, this suggests INSYN1 should be more closely investigated for its oncogenic potential.

#### Connection between the solute carrier and olfactory receptor superfamilies

One of the largest morphological clusters with novel connections (low KG scores) contains members of two superfamilies: solute carriers and olfactory receptors (OR). The cluster exists in the ORF data, not the CRISPR data (mainly because none of the olfactory receptor genes showed a phenotype after CRISPR knockout).

Among a tightly clustered subset of this large SLC-OR cluster, some gene pairs are known to be connected, but many were not (Supplementary Figures 3-5).

#### ECH1, UQCRFS1, and SARS2 cluster and are implicated in mitochondrial function and cancer

We found that three enzymes strongly correlated in both the ORF and CRISPR profiles: ECH1 (enoyl-CoA hydratase 1), UQCRFS1 (ubiquinol-cytochrome c reductase, Rieske iron-sulfur polypeptide 1), and SARS2 (seryl-tRNA synthetase 2, mitochondrial) (Figure 6 d). The connections among them are mostly unknown; UQCRFS1 and SARS2 are connected in the knowledge graph by inference through the GNN only, not by literature reports, and the remaining connections show weak or low knowledge graph scores (Figure 6 d). Some existing data supports these new connections: SARS2 is the most highly correlated gene with ECH1 in terms of cell line RNA expression; SARS2 and ECH1 are the 5th and 10th top matches, respectively, for UQCRFS1 (https://www.proteinatlas.org/ENSG00000104823-ECH1/cell+line). Analyzing the top 52 genes most similar to this cluster in the ORF dataset using Plex, the majority are mitochondria-associated (GOCC_Mitochondrion), including 16 mitochondrial disease-associated genes ^48^. Knockdown or overexpression of LINC00473, a regulator of lipolysis and mitochondrial respiration, downregulates 15 of these genes ^49^, and knockdown of the mitochondrial chaperone PHB2 downregulates 14 genes in the cluster ^50^.

Additionally, in proteomic profiling data from cells treated with a library of 875 compounds, five of the 10 profiles with the greatest overlap with this 52 gene cluster were from inhibitors of the PI3K/MTOR pathway ^51^, which regulates both mitochondrial function and biogenesis ^52^. Given recent interest in UQCRFS1 as a mitochondrial-related oncology biomarker and drug target ^53^, SARS2 and ECH1 also merit attention.

#### Connection of TSC22D1 to genes with neural function

Across both ORF and CRISPR data, we identified three strongly correlated genes (GPR176, CHRM4, TSC22D1) and a fourth gene, MYT1, consistently negatively correlated to that group (Figure 6 e). GPR176, CHRM4, and MYT1 are known to be involved in neural development or upregulated in neurons ^54,55^, and their relationship is already known, according to moderate to high scores in the knowledge graph. By contrast, TSC22D1 has little known connection to any of these and is instead annotated as involved in apoptosis, tumor suppression, and cellular stress response. The morphological similarity we observed indicates that TSC22D1 warrants further investigation for neural functions — indeed, it is most highly expressed in brain tissue according to the Human Protein Atlas (https://www.proteinatlas.org/ENSG00000102804-TSC22D1/tissue).

Furthermore, TSC22D1 is strongly anti-correlated to LZTS2 in our ORF dataset (LZTS2 was not present in our CRISPR dataset), and the two genes show an inverse mRNA expression relationship ^54,56^. LZTS2’s role in the Wnt pathway ^57^ provides another tie to neuronal function ^58^. Searching public datasets in Plex, we found that knock-down of Musashi-1 (MSI1), a gene that may play an essential role in nervous system development, in SU_MB002 medulloblastoma cells leads to down-regulation of GPR176, CHRM4, and TSC22D1 and up-regulation of MYT1 by RNA seq analysis, mirroring our observed correlations ^59^. Additionally, GPR176, CHRM4, and TSC22D1 are reported to be down-regulated in TREM2 variants associated with Alzheimer’s Disease (AD) ^60^. Together, these data indicate investigating TSC22D1 in brain function would be worthwhile.

#### Public portals for exploring JUMP data

Several portals have been built to explore and analyze the JUMP genetic perturbation data. The Broad Institute’s web-based portal JUMP Cell Painting Hub (http://broad.io/jump) allows searching for genes with the most similar profiles to a query gene, and displays images and distinctive features (relative to negative controls), to aid in interpreting the image-based phenotypes and identifying technical artifacts. Additionally, the page includes instructions for interacting with the profiles of phenotypically active ORF and CRISPR genes in the web-based data visualization and analysis software Morpheus (https://software.broadinstitute.org/morpheus/) using the compatible files stored at https://doi.org/10.5281/zenodo.14025601 and https://zenodo.org/records/14165010 and instructions at the JUMP Cell Painting Hub.

JUMP Consortium partner Ardigen developed a free web-based application, phenAID JUMP CP Data Explorer (https://phenaid.ardigen.com/jumpcpexplorer/), that enables viewing images and metadata corresponding to each genetic perturbation and searching for and downloading up to 100 nearest neighbors. By contrast to the Broad portal, whose gene-gene similarities are based on simple cosine similarities between image profiles, phenAID tool is based on projecting the ORF and CRISPR profiles to the same two-dimensional space using the UMAP algorithm with Euclidean distance and color-coded by clusters as determined using the BIRCH clustering algorithm.

## Discussion

This study presents a valuable resource for the research community: a large-scale dataset linking both over- and under-expression genetic perturbations to cell image phenotypes. We have demonstrated its utility in screening for particular phenotypes of interest and uncovering previously unknown connections between genes. The two modalities — overexpression by ORFs and underexpression by CRISPR-Cas9 knockout — provide complementary information rather than yielding identical gene clustering or simply producing opposing phenotypic profiles, consistent with our previous 160-gene study ^20^. Further investigation using this expanded dataset could provide more definitive answers, potentially through re-processing ORF and CRISPR profiles with a common pipeline to enable direct comparison rather than just at the clustering level.

This dataset has some limitations. Most notably, only 15,243 genes were tested by a reagent in the study, and only 10,352 genes had a detectable phenotype by overexpression, CRISPR, or both. The Cell Painting assay captures a broad set of phenotypes, but they are only a subset of all cell morphological changes that might occur with a broader range of cellular markers. The experiment was done on a single cell type (U-2 OS) from a single demographic (white female) at a single time point after genetic perturbation, and results may not extend to other genetic backgrounds or experimental conditions. Some individual reagents may be faulty (e.g., CRISPR guides’ off-target effects or lack of efficacy, or unexpected mutations in overexpression reagents), and, although our processing steps attempted to mitigate them, technical artifacts impact the comparison of data collected in batches and on plate layouts that induce batch and position effects ^61^. For this reason, it is crucial to examine plate layouts to ensure observed connections are not artifacts (Supplementary Figures 6-19): reagents within the same plate, row, or column can exhibit spurious correlations. Additionally, visual inspection is useful to identify image-based anomalies (Supplementary Figures 20-32).

The JUMP Cell Painting Consortium is pleased to present this genetic perturbation dataset for the scientific community’s exploration. All images and extracted profiles are freely available at https://registry.opendata.aws/cellpainting-gallery/.

## Acknowledgments

The authors thank the scientists across the entire JUMP Cell Painting Consortium for their guidance and support throughout the project.

The authors gratefully acknowledge a grant from the Massachusetts Life Sciences Center Bits to Bytes Capital Call program for funding the data production and catalyzing this Consortium. We appreciate funding to support data analysis and interpretation from members of the JUMP Cell Painting Consortium (Amgen, AstraZeneca, Bayer AG, Biogen, Eisai, Janssen Pharmaceutica NV, Merck KGaA, Darmstadt, Germany, Pfizer, Servier, Takeda Development Center Americas, Inc. (TDCA)), from the National Institutes of Health (NIH MIRA R35 GM122547 to AEC), and from grant number 2020-225720 to BAC from the Chan Zuckerberg Initiative DAF, an advised fund of the Silicon Valley Community Foundation. We would like to acknowledge the Consortium’s Supporting Partners for their contributions: Ardigen for their deep learning expertise and JUMP-CP Data Explorer web application (part of Ardigen’s phenAID platform); Google/Verily for the compute support and configuration/optimization of Terra, which is co-developed by the Broad Institute of MIT and Harvard, Microsoft and Verily (its use is not described in this paper); Nomic bio for their protein profiling (not described in this paper); and Revvity for the PhenoVue^TM^ Cell Painting JUMP kit and the Edit-R Libraries and Edit-R tracrRNA. We also are grateful for the Amazon Web Services Registry of Open Data for hosting the public dataset. The authors also acknowledge the use of the Revvity Opera Phenix^TM^ High-Content/High-Throughput imaging system at the Broad Institute, funded by the S10 Grant NIH OD-026839.

## Authors’ contributions

**Srinivas Niranj Chandrasekaran**: Analysis, Data curation, Experiments/Investigation, Methodology, Project administration, Software, Supervision, Validation, Visualization, Writing – original draft, Writing – review & editing; **Eric Alix**: Experiments/Investigation, Resources; **John Arevalo**: Analysis, Data curation, Software, Validation; **Adriana Borowa**: Methodology, Supervision, Writing – original draft; **Patrick J. Byrne**: Data curation, Experiments/Investigation, Validation; **William G. Charles**: Data curation, Software, Visualization; **Zitong S. Chen**: Analysis, Data curation, Experiments/Investigation, Methodology, Software, Validation, Writing – review & editing; **Beth A. Cimini**: Data curation, Methodology, Project administration, Software, Supervision, Writing – original draft, Writing – review & editing; **Boxiong Deng**: Experiments/Investigation; **John G. Doench**: Project administration, Resources, Supervision; **Jessica D. Ewald**: Experiments/Investigation, Writing – original draft; **Briana Fritchman**: Experiments/Investigation, Methodology, Project administration, Resources, Supervision, Writing – original draft; **Colin J. Fuller**: Analysis, Data curation, Software, Validation, Writing – review & editing; **Jedidiah Gaetz**: Analysis, Writing – original draft; **Amy Goodale**: Experiments/Investigation, Methodology, Project administration, Resources, Supervision, Writing – original draft; **Marzieh Haghighi**: Analysis, Experiments/Investigation, Software; **Yu Han**: Analysis, Software, Validation; **Zahra Hanifehlou**: Analysis, Data curation, Validation; **Holger Hennig**: Analysis, Conceptualization, Methodology, Project administration, Supervision, Visualization, Writing – original draft; **Desiree Hernandez**: Experiments/Investigation, Resources; **Christina B. Jacob**: Data curation, Experiments/Investigation, Project administration, Supervision; **Tim James**: Funding acquisition, Supervision; **Tomasz Jetka**: Analysis, Methodology, Project administration, Supervision, Visualization, Writing – original draft; **Alexandr A. Kalinin**: Analysis, Experiments/Investigation, Methodology, Software, Writing – review & editing; **Ben Komalo**: Analysis, Data curation, Software, Validation, Writing – review & editing; **Maria Kost-Alimova**: Data curation, Experiments/Investigation, Methodology, Project administration, Resources, Supervision, Writing – review & editing; **Tomasz Krawiec**: Software, Supervision; **Brittany A. Marion**: Experiments/Investigation, Methodology, Project administration, Resources, Validation; **Glynn Martin**: Experiments/Investigation, Methodology, Resources; **Nicola Jane McCarthy**: Experiments/Investigation, Funding acquisition, Supervision; **Lisa Miller**: Experiments/Investigation, Methodology, Project administration, Validation, Writing – original draft; **Arne Monsees**: Analysis, Data curation, Methodology, Software, Visualization, Writing – original draft, Writing – review & editing; **Nikita Moshkov**: Analysis; **Alán F. Muñoz**: Analysis, Data curation, Experiments/Investigation, Methodology, Software, Validation, Writing – original draft, Writing – review & editing; **Arnaud Ogier**: Analysis, Project administration, Supervision; **Magdalena Otrocka**: Supervision, Writing – original draft, Writing – review & editing; **Krzysztof Rataj**: Analysis, Data curation; **David E. Root**: Methodology, Project administration, Resources, Supervision; **Francesco Rubbo**: Analysis, Data curation, Software, Validation, Writing – review & editing; **Simon Scrace**: Experiments/Investigation, Project administration, Resources; **Douglas W. Selinger**: Analysis, Writing – original draft; **Rebecca A. Senft**: Analysis; **Peter Sommer**: Conceptualization, Supervision; **Amandine Thibaudeau**: Experiments/Investigation, Validation; **Sarah Trisorus**: Analysis, Data curation, Software, Validation, Writing – review & editing; **Rahul Valiya Veettil**: Analysis, Data curation, Methodology, Software, Visualization, Writing – original draft, Writing – review & editing; **William J. Van Trump**: Analysis, Data curation, Methodology, Software, Supervision, Visualization, Writing – original draft, Writing – review & editing; **Sui Wang**: Analysis, Data curation, Experiments/Investigation, Validation, Writing – review & editing; **Michał Warchoł**: Project administration, Supervision, Writing – review & editing; **Erin Weisbart**: Analysis, Data curation, Experiments/Investigation, Software, Validation, Writing – review & editing; **Amélie Weiss**: Experiments/Investigation, Validation; **Michael Wiest**: Analysis, Data curation, Software, Validation, Writing – review & editing; **Agata Zaremba**: Analysis, Visualization; **Andrei Zinovyev**: Analysis, Data curation, Methodology, Software, Supervision, Visualization, Writing – original draft, Writing – review & editing; **Shantanu Singh**: Analysis, Conceptualization, Data curation, Experiments/Investigation, Funding acquisition, Methodology, Project administration, Software, Supervision, Validation, Visualization, Writing – original draft, Writing – review & editing; **Anne E. Carpenter**: Analysis, Conceptualization, Funding acquisition, Methodology, Project administration, Supervision, Visualization, Writing – original draft, Writing – review & editing

## Data and code availability

The data is available for free download from the Cell Painting Gallery AWS S3 bucket without requiring an AWS account (accession number: cpg0016). Detailed instructions for downloading the data are available at https://github.com/jump-cellpainting/2024_Chandrasekaran_Morphmap/blob/main/README.md. The code for generating all the figures is available in our GitHub repository: https://github.com/jump-cellpainting/2024_Chandrasekaran_Morphmap

## Methods

### Genetic perturbation selection

#### ORF expression library

For cpg0016, we used a preexisting lentiviral ORF expression library created at the Broad Institute for gene overexpression experiments ^23^. We tested 15,141 overexpression reagents encompassing 12,609 unique genes, including controls, which are described in a later section. Lentiviral packaging becomes less efficient for larger genes; this library does not cover the entire genome because it excludes larger genes.

CPJUMP1 data (cpg0000) also contains ORF perturbation data, where we overexpressed 176 genes. Additional details about these perturbations have been published ^20^.

#### CRISPR knockout library

We used the Revvity Human Edit-R^TM^ synthetic crRNA - Druggable Genome in our CRISPR knockout experiments. This library targets 7,975 unique genes; each targeted by a pool of four predesigned synthetic crRNAs with high specificity and functional knockout.

In the CRISPR knockout experiments in CPJUMP1, we knocked down 160 genes. Additional details about these perturbations have been published ^20^.

#### Controls

We included several controls to identify and correct for different experimental artifacts. The controls can be broadly categorized as within-plate controls (those on the same plate as the treatments) and control plates (entire plates run alongside the treatment plates).

#### Within-plate controls

Within-plate controls include negative controls, positive controls, and untreated wells.

Negative control wells: These controls are used to detect and correct plate-to-plate variations. They can also be used as a baseline for identifying perturbations with a detectable morphological signal. The following are the within-plate negative controls.

● ORF plates – ORFs of genes BFP, HcRed, lacZ, and Luciferase, although it should be noted that no sensible negative control exists for protein overexpression, as each of these proteins is known to induce some changes to cell state.
● CRISPR plates – Non-targeting guides (CRISPR guides that do not target any gene) and no guides (Cas9 cells are transfected but did not receive any CRISPR guides). There are also wells where cells are treated with DMSO but are not used as negative control treatments.

Positive control wells: These controls are included in the plates to ensure the experiment worked as expected.

● ORF plates – We ran four positive controls on the ORF plates (Supplementary Table 3) which are a subset of the eight positive control compounds run on the CRISPR plates. Furthermore, we included eGFP (enhanced Green Fluorescent Protein) as the ORF positive control, which showed a consistent phenotype distinct from the negative controls.
● CRISPR plates – PLK1; knocking it down kills cells. We also ran eight compound positive controls on each CRISPR plate (Supplementary Table 3). These compounds exhibited phenotypes that are distinct from each other and from the negative control in previous experiments ^20^.

Untreated wells: Some wells received neither treatments nor controls; they contained only cells. They might also be used for normalizing samples, but they are non-ideal negative controls, given that they are not mock-treated with any reagents.

#### Control plates

Negative control plates: Negative control plates we run periodically (e.g., one or more per batch) (Figure 1 b and c). These plates can help disentangle staining and imaging artifacts, including batch effects and well-position effects. Examples include:

● An entire plate of untreated cells.
● An entire plate of cells treated with non-targeting or no guides (for CRISPR).

JUMP-Target-2-Compound plates: One or more plates of JUMP-Target-2-Compound compounds were run in each batch to correct batch effects (unrelated plates from the same batch correlate strongly with each other compared to related plates in other batches)(Figure 1 c).

JUMP-Target-1-Compound plates: To align the production data (cpg0016) with the CPJUMP1 experiment in the pilot data (cpg0000) ^20^, we ran four JUMP-Target-1-Compound plates in one batch of cpg0016 from source_4 (Broad) (Figure 1 b and c).

JUMP-Target-2-Compound plates with polybrene: To quantify the cell morphology effects of adding polybrene, a viral transfection agent that is added to the ORF experiment, we ran two JUMP-Target-2-Compound plates with polybrene in one batch (Figure 1 b and c).

### Plate layout design

#### ORF plates

The ORF plates were pre-designed due to their existence in a pre-plated library ^23^; each plate consists of negative control wells and untreated wells spread across the plate. Most plates contain 16 negative control wells, while some have as many as 28 wells. One replicate for each of the four compound positive controls (Supplementary Table 3) is added to wells O23, O24, P23, and P24. The remaining wells contain ORF treatments, with a single replicate of each per plate map and with five replicate plates produced per plate map.

#### CRISPR plates

Similar to the compound plates, the outer columns of the plate contain positive and negative controls. The outermost columns 1 and 24 contain four replicates of the eight compound positive controls. Columns 2 and 23 contain ten replicates each of wells with no guides and non-targeting guides, eight replicates of DMSO, and four replicates of the CRISPR positive control, PLK1.

#### JUMP-Target-1-Compound plates

The plates contain 306 compounds and DMSO; all but 14 compounds are in singlicates. There are 64 DMSO wells spread across the plate. The 14 compounds with two replicates are diverse positive controls that differ from the positive controls on the production plate. Additional details about these positive controls and the criteria met by the plate’s compounds have been described ^20^.

#### JUMP-Target-2-Compound plates

These plates’ contents are identical to those of JUMP-Target-1-Compound plates, but the layout differs.

JUMP-Target-2-Compound plates meet all the criteria met by JUMP-Target-1-Compound plates ^22^. Both layouts are provided in our GitHub repository (https://github.com/jump-cellpainting/JUMP-Target).

### Perturbation identifier

All ORF reagents, CRISPR reagents, and controls are assigned a unique JUMP identifier. These identifiers start with the code *JCP2022_*.

### Experimental conditions

#### Cell Painting dyes

We optimized the concentrations of several Cell Painting dyes and chose the concentration as described in our previous publication ^22^.

#### Cell line

After comparing A549 and U-2 OS cell lines in pilot experiments ^20^, we selected U-2 OS because of its performance in Cell Painting experiments and the existence of previous datasets in this cell line, allowing comparison across experiments.

#### Time point

We compared two time points for each perturbation modality: 48 and 96 hours of ORF and 96 and 144 hours for CRISPR and settled on 48 hours for ORF and 96 hours for CRISPR, based on their performance ^20^.

#### Reagent vendor

Cell Painting dyes from Thermo Fisher and Revvity performed similarly in our pilot experiments ^22^. For this dataset, we used the PhenoVue^TM^ Cell Painting Kit, 2.0 (part number PING22, Revvity, Waltham, MA), which Revvity donated to the JUMP Cell Painting consortium.

#### Microtiter plates

After comparing different plates for their ability to minimize evaporation in the outer wells (cpg0000) ^22^, we used Revvity Cell Carrier Ultra for data generation.

#### Compound concentration

The positive control compounds in the ORF and CRISPR plates, JUMP-Target-1-Compound and JUMP-Target-2-Compound plates, were assayed at 5 μM.

#### ORF experiments

We followed previous protocols ^62,63^, with the following experimental parameters chosen: 5 replicates per virus plate, 1525 cells/384w, 30 μL media seeding volume/384w, 1 μL virus/384w, and 4 μg/mL polybrene added 1 hour post-seeding, 30-minute spin at ∼1000g, media change after 24 hrs removing polybrene and virus adding back 40 μL media, no selection with Blasticidin.

#### Creation of the Cas9 cell line

We made U2OS-311 at the Broad Institute by transducing U-2 OS cells at a multiplicity of infection (MOI) <1 with lentivirus prepared from the vector pLX_311-Cas9 (Addgene plasmid 96924), which expresses blasticidin resistance from the SV40 promoter and Cas9 from the EF1α promoter, and selected with 16 μg/ml blasticidin for 14 d. Briefly, a 12-well plate at 1.5 × 10e6 cells per well in 1.25 mL media and 750 μL virus supplemented with 4 μg/ml polybrene were centrifuged for 2 h at 1,000*g,* then 2 mL media was added per well. 24 h after infection, cells were split out of the 12-well plate, and 48 h after infection, 16 μg/ml blasticidin was added and maintained. This is a slight modification of our published protocol ^64^.

#### CRISPR experiments

Five replicates per target gene were assessed, with each replicate well containing four different sequences targeting the same gene. The cell line used for these experiments was U2OS-311 (Broad). The cells were kept for a week in blasticidin before plating. Blasticidin was removed one passage before plating. Before plating, 125nL of each guide RNA pool at 10μM (Revvity) and 125nL of the tracrRNA at 10μM (Revvity) were dispensed in the 384 well plates using the ECHO (Beckman Coulter Life Science). 10uL of LipoRNAimax (Thermo Fisher) diluted in OptiMEM was added to each well using a MultiDrop Combi (Thermo Fisher) to achieve a final 0.05uL/well of LipoRNAimax. The plates were pulse centrifuged and then incubated for 30 minutes at room temperature. Then 700 cells per well were dispensed in 40uL using the MultiDrop Combi and incubated for 96h in an incubator (37C, 5% CO2) before staining and fixation. Additional chemical treatments were included in some control wells (DMSO, C1, C2, … C8), and some full plates of cells were treated with the Target-2 plate of compounds. These treatments were performed as described in the chemical screening and were incubated for 48h before staining and fixing U2OS-311 cells with LipoRNAimax treatment only (no guide RNA, no tracrRNA).

### Sample preparation and image acquisition

The Cell Painting assay employs six fluorescent dyes, imaged across five channels, to visualize eight key cellular components: mitochondria (MitoTracker, Mito), nucleus (Hoechst, DNA), nucleoli and cytoplasmic RNA (SYTO 14, RNA), endoplasmic reticulum (concanavalin A, ER), Golgi and plasma membrane (wheat germ agglutinin) plus the actin cytoskeleton (phalloidin, AGP). We followed the optimized Cell Painting assay protocol ^22^ (https://github.com/carpenterlab/2022_Cimini_NatureProtocols/wiki). Image acquisition for the ORF dataset was performed using the Revvity Opera Phenix microscope in the widefield mode and for the CRISPR dataset using Yokogawa CV7000 in the confocal mode. Only fluorescent channels were acquired for the CRISPR dataset, while three brightfield planes were also acquired for the ORF dataset. Nine fields of view were acquired for both the ORF and CRISPR datasets.

### Image Processing

We used CellProfiler bioimage analysis software (versions 4.1.3 or 4.2.1) for image processing. We corrected background illumination variations ^65^ and segmented nuclei and cells using classical segmentation algorithms, namely, CellProfiler’s IdentifyPrimaryObjects module with Minimum Cross-Entropy thresholding and IdentifySecondaryObjects module with Otsu three-class watershed thresholding, respectively. Subsequently, we measured a suite of features for each cell in every fluorescent channel, including intensity, granularity, texture, and density. We extended this analysis to brightfield planes for the ORF dataset. Additionally, we quantified features at the whole-image level. (Refer to http://broad.io/cellprofilermanual for detailed methodology. Our image analysis pipeline (see https://github.com/broadinstitute/imaging-platform-pipelines/tree/fc10d6acec5c1b2d9c4526a052acb1d9a196525a/JUMP_production#production-pipelines) generated up to 7648 features (encompassing both per-cell and per-image measurements) for the ORF dataset. For the CRISPR dataset, we extracted 4762 features per cell, as we didn’t measure any brightfield features.

### Quality control

We implemented several quality control measures to improve data quality. First, we excluded Batch 12 from the ORF dataset due to incorrect dispensing of assay dyes in two plate rows. This batch was subsequently repeated as Batch 13. Next, we filtered ORF reagents based on viral infection efficiency, as determined by a CellTiter-Glo® cell viability assay conducted in parallel with the Cell Painting plates. The infection efficiency values showed a bimodal distribution, with two outlier reagents (Supplementary Figure 33). We established a threshold for low infection efficiency using Otsu thresholding. This filtration process eliminated all EMPTY wells, as expected, due to the absence of virus in these wells. The unique, non-symmetrical layout of EMPTY wells in each plate map confirmed that plates were neither swapped nor rotated relative to their metadata. One exception was noted: plate map OAB41.OAC17.OAB78.79.A in Batch 04, where three EMPTY wells displayed strong infection efficiency. As the pattern did not improve with rotation and lacked a plausible explanation, we opted to exclude all five replicates of this plate map from the experiment. We also removed control wells and ORF reagents with infection efficiency below the Otsu threshold. Together, these quality control measures resulted in the removal of 2,397 ORF reagents. The infection efficiency of each ORF reagent is available at https://github.com/jump-cellpainting/2024_Chandrasekaran_Morphmap/blob/0ba4e4bdd4ee18cf6dae20d4af48076a53c1d47f/00.download-and-process-annotations/input/JUMP-ORF-Infection-Efficiencies.xlsx and the list of removed ORF reagents is available at https://github.com/jump-cellpainting/2024_Chandrasekaran_Morphmap/blob/3fdd5e32bde0ae51dd6cbab6d658fe11667806c0/00.download-and-process-annotations/output/orf-reagents-low-infection-efficiency-and-outliers.csv.gz.

### Overview of profile processing workflow

We extracted single cell features with CellProfiler, following steps described in the profiling handbook ^66^. Then, we mean-aggregated the profiles using Pycytominer ^67^ to generate well-level profiles. To correct for technical artifacts in the dataset, we began by correcting for systematic variations across well positions. We calculated the mean value for each feature within each well position and subtracted this mean from the corresponding feature values of individual samples. Next, recognizing that cell count can strongly contribute to data variability ^16^, we addressed its influence on other features by regressing it out while retaining the original cell count feature for subsequent analyses. Following this, we applied a modified version of our profile processing pipeline for batch correction ^61^. We applied variance thresholding to remove features with minimal variation across the dataset. We then scaled the data based on its median absolute deviation and performed feature selection to remove invariant and redundant features. Finally, we performed sphering transformation to decorrelate features and standardize variance, followed by Harmony batch correction ^68^ to mitigate technical artifacts while preserving biological variation.

CRISPR-Cas9-mediated gene knockouts, while intended to disrupt specific genes through targeted DNA cuts, can occasionally result in unintended large-scale deletions encompassing the entire remaining chromosome arm ^69–71^. Recent studies have identified systematic patterns in image-based and cell-line viability profiles, revealing similarities among knockouts of genes located on the same chromosome arm ^19^. We confirmed that this pattern is seen in our CRISPR dataset (https://doi.org/10.5281/zenodo.13754407), which had undergone different pre-processing than in the prior study, but, as expected, not in our overexpression dataset (https://doi.org/10.5281/zenodo.13754178). The proposed correction method ^19^, involving PCA followed by correction, successfully mitigated this pattern (https://doi.org/10.5281/zenodo.13754508). This correction was applied only to the CRISPR dataset.

This processing workflow can be replicated by running https://github.com/broadinstitute/jump-profiling-recipe/tree/v0.1.0 using the appropriate config file for the CRISPR (crispr.json) and ORF datasets (orf.json).

Mean average precision (mAP) calculations for phenotypic activity and phenotypic consistency are detailed and defined ^24^. For example, mAP measures phenotypic activity when we calculate each replicate’s ability to retrieve the other replicates for that gene against the background of negative control samples, using cosine similarity as the similarity metric. We assess statistical significance using permutation testing to obtain a p value. P values are adjusted for false discovery rate to yield a corrected p value (q value).

### Gene annotations

Annotations were downloaded from publicly available databases as described in

https://github.com/jump-cellpainting/2024_Chandrasekaran_Morphmap/blob/5e6da3cae56443584a89cd9205a 2e4190d8ebbaf/00.download-and-process-annotations/README.md. To transform the predominantly continuous annotation data into categorical form for the Fisher’s exact test, we established threshold values.

For essentiality, we set the dependency probability threshold as 0.75, and for expression, we set the TPM threshold as 75. For ORF gene insert size, we set the threshold as 2500 nucleotides.

### Single Cell PhenoSorter Models

Raw images were uploaded to Spring Engine (a cloud-based software-as-a-service product provided by Spring Science). We used the PhenoSorter application to generate a single-cell-based phenotypic model that discriminated between flat and rounded cells. Briefly, this application presents individual segmented cell crops to a user, who classifies them as rounded cells or flat cells. These cell crops are processed using RepLKNet, a pre-trained large kernel convolutional neural network ^72^. The final layer is pooled using average pooling to yield a 128-element embedding vector for each imaging channel. Classification models are built upon this embedding vector using XGBoost ^73^, a gradient-boosting decision tree-based framework. For this model, embeddings were concatenated from all five fluorescence channels used in the ORF dataset. The training uses an 80/20 split in which 80% of the user-classified cell images are randomly chosen to train the model, and the remaining 20% are tested to determine model performance. To accommodate for different example set sizes, the loss function is weighted so that examples from smaller classes are given greater weight, such that each class is ultimately balanced despite the different sample set sizes. Once trained, the model is applied to all single-cell image crops in the entire dataset. The data for all cells in each well are aggregated to yield a percent value for each model class for every well, and we applied a threshold of 6.5% reduction in the percentage of living cells compared to negative controls to call hits.

### Gene Set Enrichment Analysis for PhenoSorter flat/rounded model

We evaluated the results of our ML classification models by performing a gene set enrichment analysis on the ORF perturbations that demonstrated the greatest effect of the model. For the rounded cell model, we compared the fraction of cells within each well that the model scored as flat for each ORF treatment against that same fraction of cells for untreated cells. ORF treatments were then ranked by the percent reduction in flat cells. A cutoff of 6.5% increase in rounded cells was used to generate our gene subset for genes demonstrating increased levels of cell roundedness. The R programming language package ‘clusterProfiler’ ^74^ was used for gene set enrichment analysis. The WikiPathways annotation set of the C2 subset of annotations sourced from the MSig database was used for gene annotations.

### Identifying relevant datasets using Plex

Genes sets in selected clusters (i.e. genes similar to YAP1 in the ORF data, genes positively or negatively correlated with TSC22D1, and the cluster containing ECH1, UQCRFS1, and SARS2) were searched in the Plex Research web app (https://plexresearch.com). Plex combines knowledge graphs with centrality algorithms to find convergences between search inputs and diverse, large, publicly available datasets such as Gene Expression Omnibus^75^, Opentargets (https://www.opentargets.org), the Broad Cancer Dependency Map Project (https://depmap.org/portal).

### Gene-gene functional connections quantified from a knowledge graph

Biomedical knowledge graphs (KG) represent information in the form of nodes (also called entities) connected by edges (also called links), where entities can have various types (e.g., gene, compound, biological function, pathway, disease, etc.) and the links can represent various facts connecting them (e.g., a gene involved in a pathway or a disease, two genes co-cited in an article, a compound is used to treat a disease). We used the Drug Repurposing Knowledge Graph (DRKG) ^26^, a comprehensive open-source biomedical knowledge graph relating genes, compounds, diseases, biological processes, side effects, and symptoms. DRKG includes information from six existing databases, including DrugBank, Hetionet, GNBR, String, IntAct, and DGIdb, with 4,078,154 edges (of 107 edge types) and 66,500 nodes (of 13 node types), after filtering out non-human genes and edges. The DRKG graph includes the results of large-scale scientific literature mining coming from the PubTator project through the global network of biomedical relationships (GNBR).

We predict links between genes and functions in DRKG with a Graph Neural Network (GNN) approach. The analysis is limited to the 6,787 genes (ORF data) and 5,494 genes (CRISPR data) having a phenotype and existing in DRKG (the majority: 6787/7031=97%, and 5494/5546=99%). Graph representation learning methods generate vector representations for graph nodes such that the learned representations, i.e., embeddings, capture the structure and semantics of networks ^76^. To introduce a numerical score reflecting biological functional proximity between two genes from multiple sources of evidence, we trained three predictive models using the DRKG and GNN approach, aimed at predicting either molecular function of a gene using Gene Ontology (GO) Molecular Function (MF) definitions as training set, or the biological process in which a gene is involved using GO Biological Process (BP) definitions as training set, or the pathway in which a gene is involved (using definition of pathways from various resources, including WikiPathways and Reactome as training set). As an architecture for GNN, we implemented a variational graph autoencoder (VGAE) ^77^ in Pytorch Geometric ^78^ for predicting links between genes and functions, using the whole topographical structure of the knowledge graph inspired by https://colab.research.google.com/drive/1Jv0GrF11jcbhiV7dK-RhxKCv_GLVvTls?usp=sharing#scrollTo=sBWeMphti_y2. This model makes use of latent variables and is capable of learning interpretable latent representations for undirected graphs. For training, we used a standard validation strategy by retaining a part of the gene-function links from the training set as a validation set and measuring the performance of the prediction through the metrics Area Under the Curve (AUC) and Hits at k (Hits@k) ^79^. Hits@K measures the proportion of correct predictions within the top k-ranked candidates, i.e., it denotes the ratio of the test triples that have been ranked among the top k triples ^80^.

For a final validation of the models, we applied a time machine approach ^81^, where computational models are trained using data published before a certain time point, and the model outputs are validated by their ability to predict links that became part of biomedical knowledge after this time point. Specifically, for validation we used those links from the latest GO ^82,83^ that were established after 2020 and thus are not a part of DRKG, which was created in 2020. The prediction accuracy was high in both validation and test sets (Supplementary Table 4).

Each of the trained models provides a score *x_g,f_* for any pair of a gene *g* and a node *f* in DRKG representing a biological function (either GO MF category, GO BP category, or a pathway), where the known links typically score on top while other relatively large scores represent predictions of a gene function from the whole content of the knowledge graph. For a given type of biological function, the functional KG score between two genes *g_i_* and *g_j_* is computed as

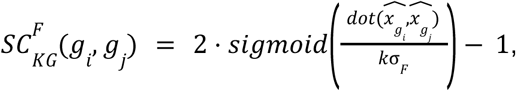

where *dot*() is the standard dot product; 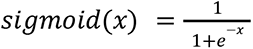 is a sigmoid function; *F* is the type of the biological function node in KG, *F*∈{GO_MF, GO_BP, pathway}; 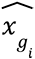 is the centered vector of scores 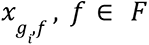; σ_*F*_ is the standard deviation of the distribution of 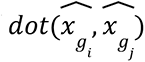 values for all possible gene pairs *g_i_*, *g_j_*; *k* is the spreading coefficient empirically chosen at 1.5 in all experiments to avoid the saturation of sigmoid function values close to 1.0.

Therefore, the functional score between two genes is a value in [-1;1] interval, with values close to 1.0 representing two closely functionally related genes (these are usually the genes involved in the same family, with physically interacting gene products, known to be involved in the same biological function, etc.) and -1.0 representing genes maximally separated in the knowledge graph. Empirically, we observed that the distribution of 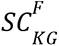 scores has the maximum density for the score close to 0.2 (see Figure 3) with a relatively small number of negative values. We use 0.5 as a cutoff for whether a particular relationship is previously known.

