## Supplementary materials for "Morphological map of under- and over-expression of genes in human cells"

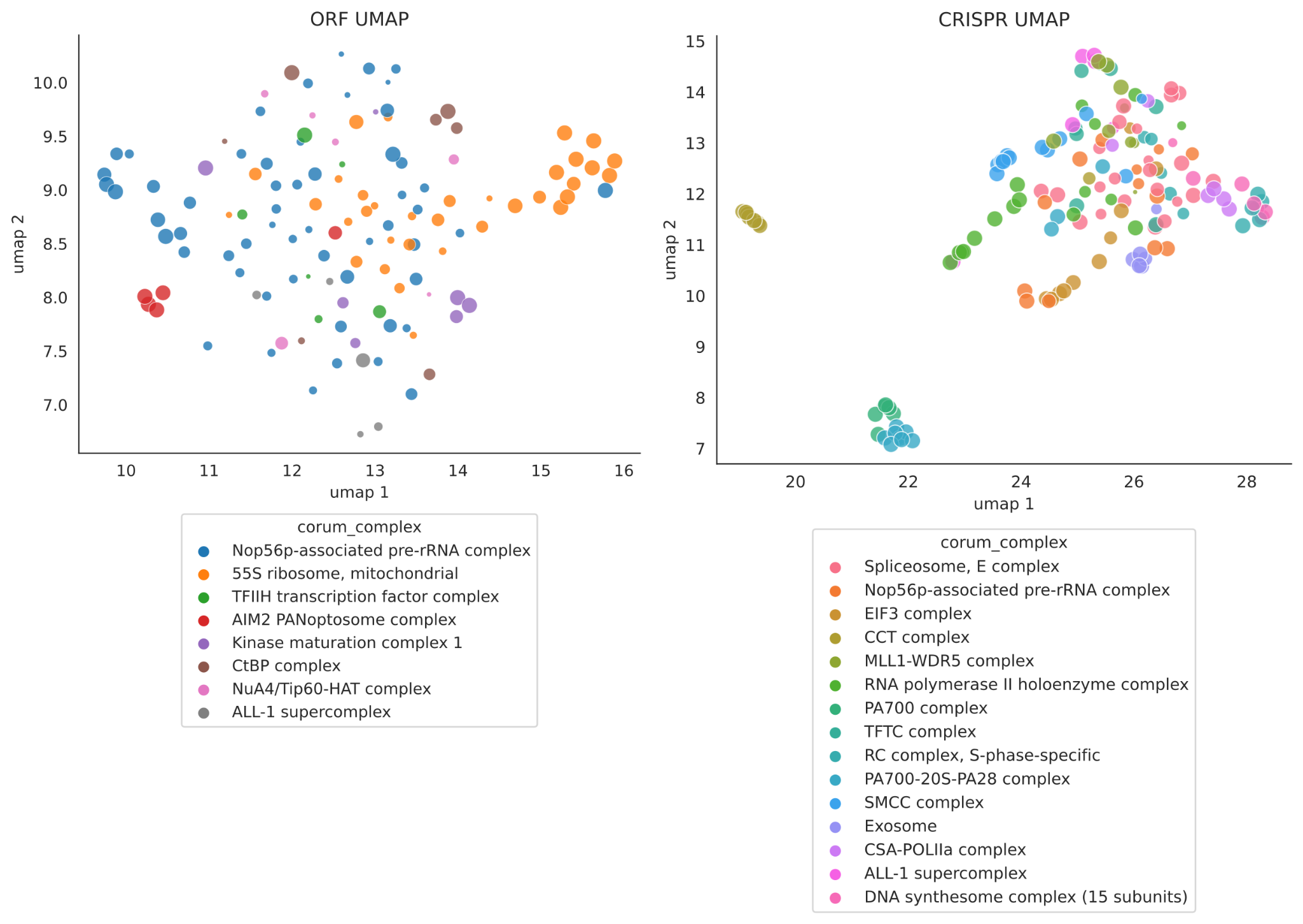

*Supplementary Figure 1****: UMAP visualization of genes based on their ORF and CRISPR Cell Painting profiles.*** *The genes shown are only those in a CORUM complex (indicated by color; size indicates the phenotypic activity mAP) with a minimum mAP of 0.1 for phenotypic consistency and at least five genes. These clusters show a strong phenotypic consistency.*

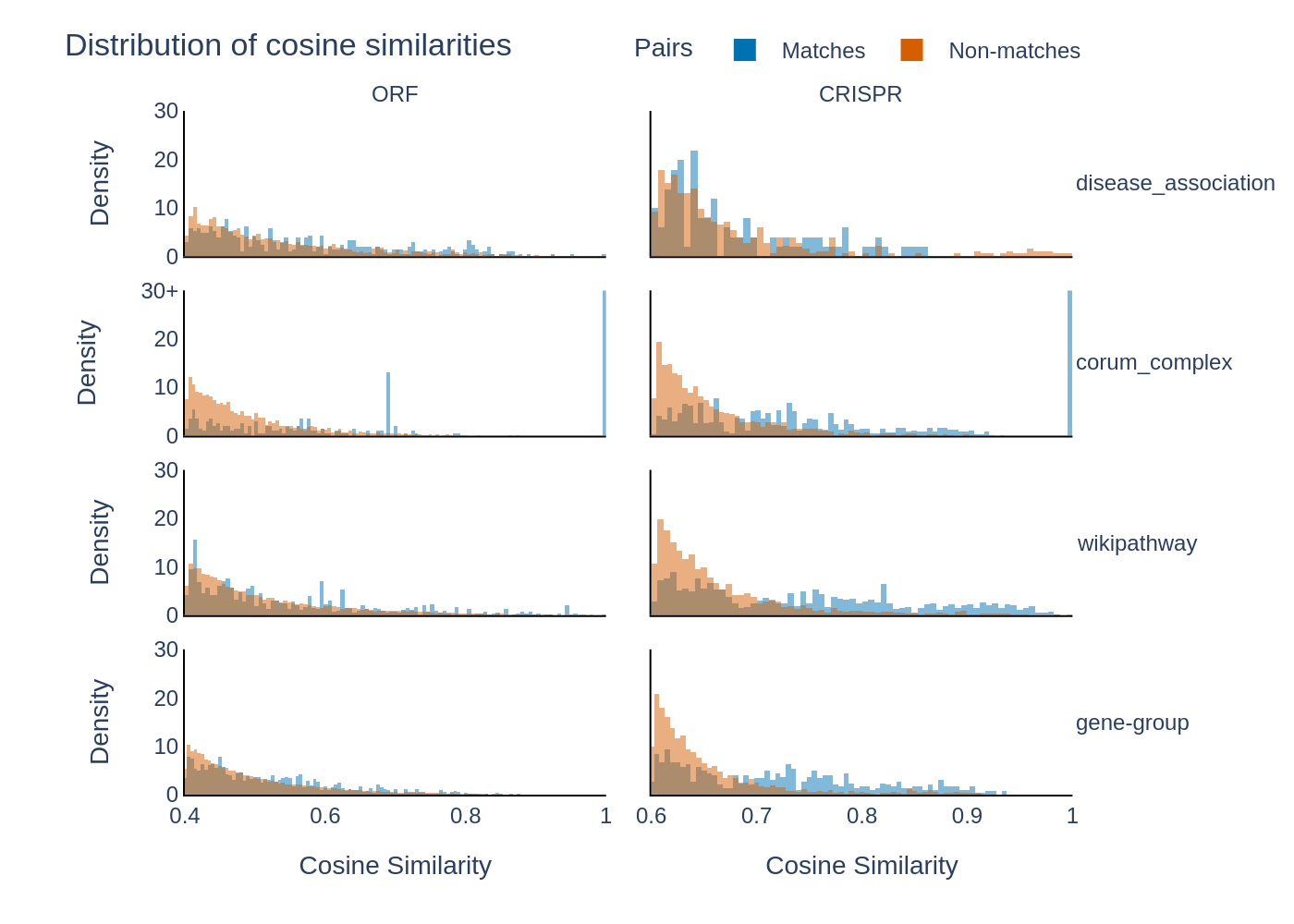

*Supplementary Figure 2**:* ***Histogram of Cosine similarities.*** *Cosine similarity of profiles of gene pairs that are in the same (blue) or different (orange) group, based on existing annotations. Only the tails of each distribution are shown here.*

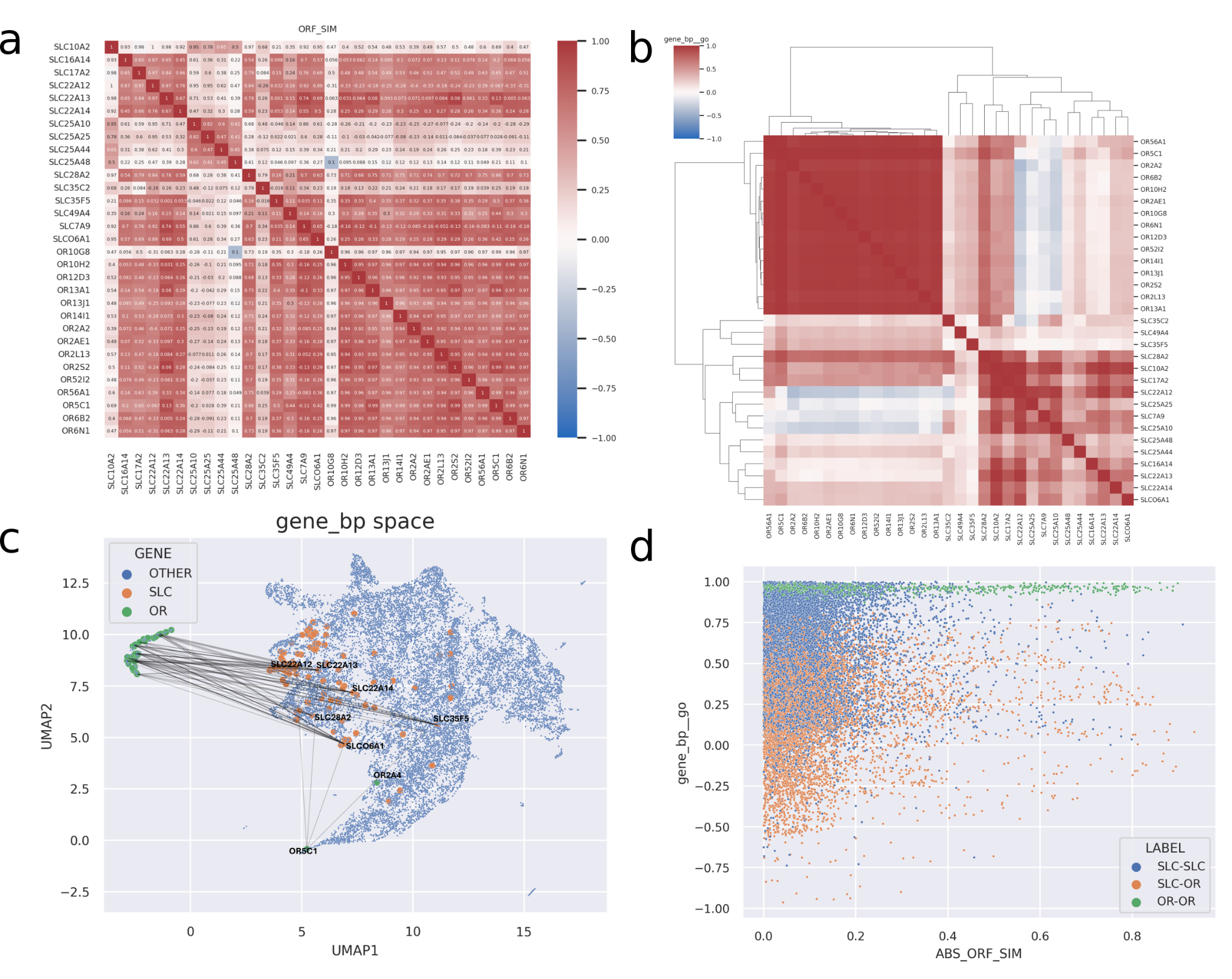

*Supplementary Figure 3**:* ***Connecting Solute Carriers and Olfactory Receptors.*** *Some members of the solute carrier (SLC) and olfactory receptor (OR) superfamilies are connected via strong ORF similarities, but these two superfamilies are relatively separated in the knowledge graph without strong evidence of the biological connection between them. a) Strong ORF similarities (larger than 0.7) between SLC and OR members (alphabetical order). The cell annotations in the clustergram indicate the KG similarity score between two genes. b) Clustering of the same genes as in A) but according to the KG similarity score. c) UMAP visualization of the KG functional embedding space. Most OR genes are isolated from other genes in the embedding (as genes about which little is known), while many of them are connected to solute carriers by ORF Cell Painting similarities. The line shows only Cell Painting ORF-defined connections between SLC and OR genes. d) Scatter plot visualizing the relation between the absolute value of ORF similarity score and the KG similarity score. It is clear that most SCL-OR links having strong ORF similarity cannot be explained from KG (they have relatively small KG similarity score).*

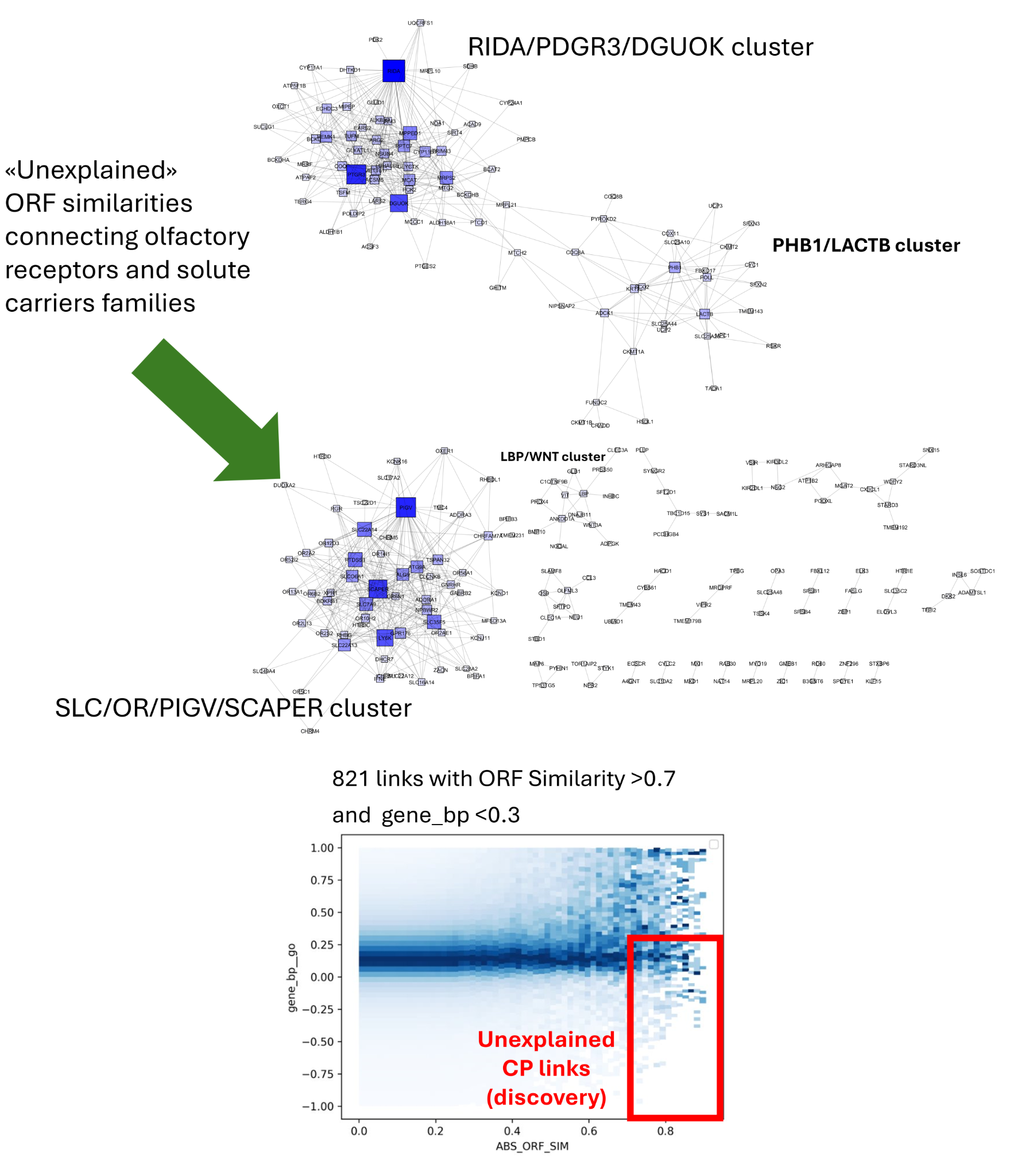

*Supplementary Figure 4**:* ***Unexplained clusters in the ORF dataset.*** *Links between genes characterized by strong ORF similarity (ABS(ORF)>0.7) that cannot be explained from the knowledge graph (KG similarity score<0.3) can be grouped into clusters of links. One of the largest clusters connects solute carriers and olfactory receptors. This and other clusters are annotated by the names of the most connected genes. Strongly connected nodes (having many similar and unexplained ORF profiles) are highlighted by size and color.*

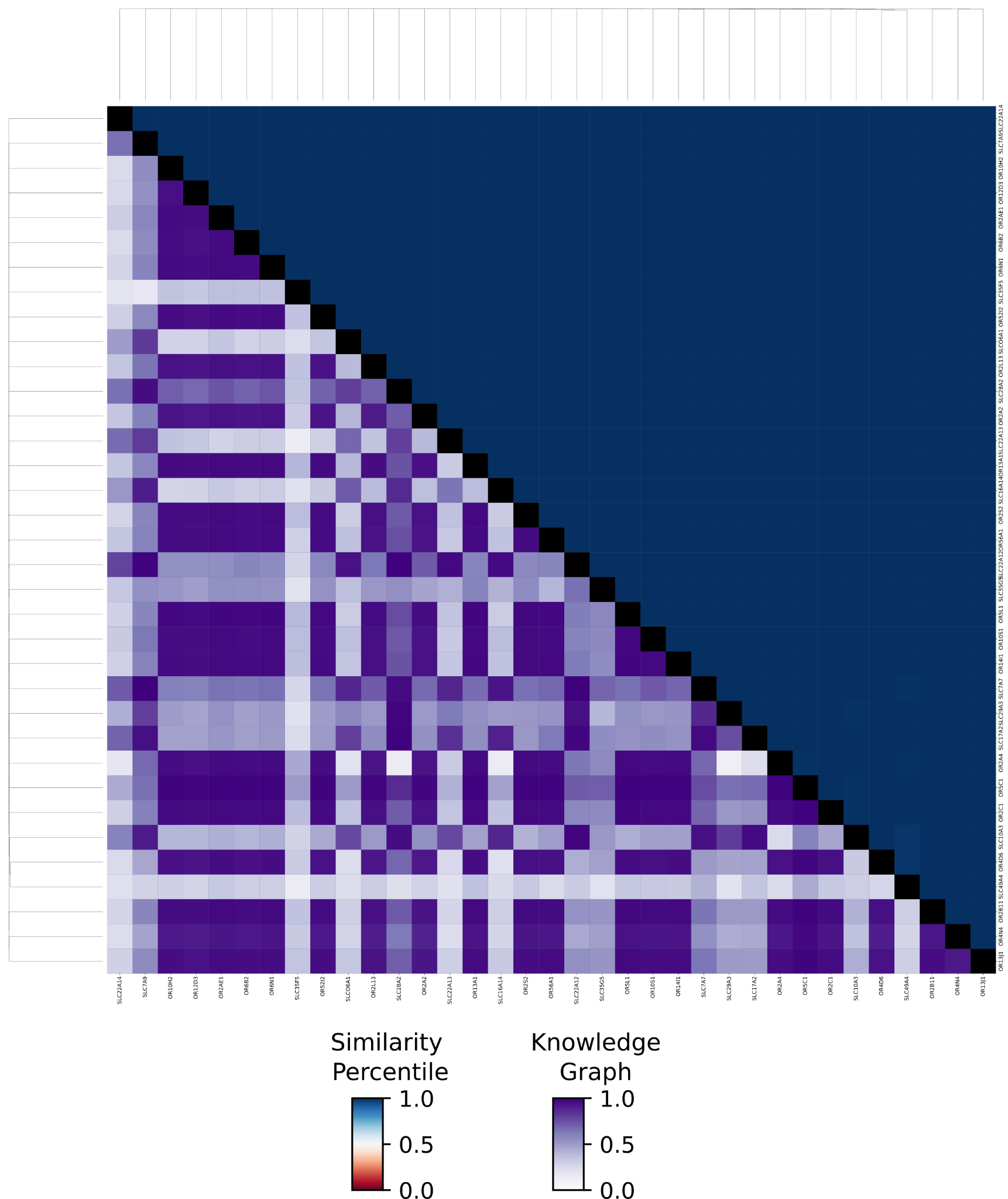

*Supplementary Figure 5**:* ***Clustergram of the tight cluster made of a subset of SLC and OR genes.*** *The upper triangular matrix displays the percentile of cosine similarity between gene profiles (0 = negative correlation, 1 = positive correlation). The lower triangular matrix shows the knowledge graph score (see Methods), with scores < 0.4 indicating novel associations not previously reported in the literature.*

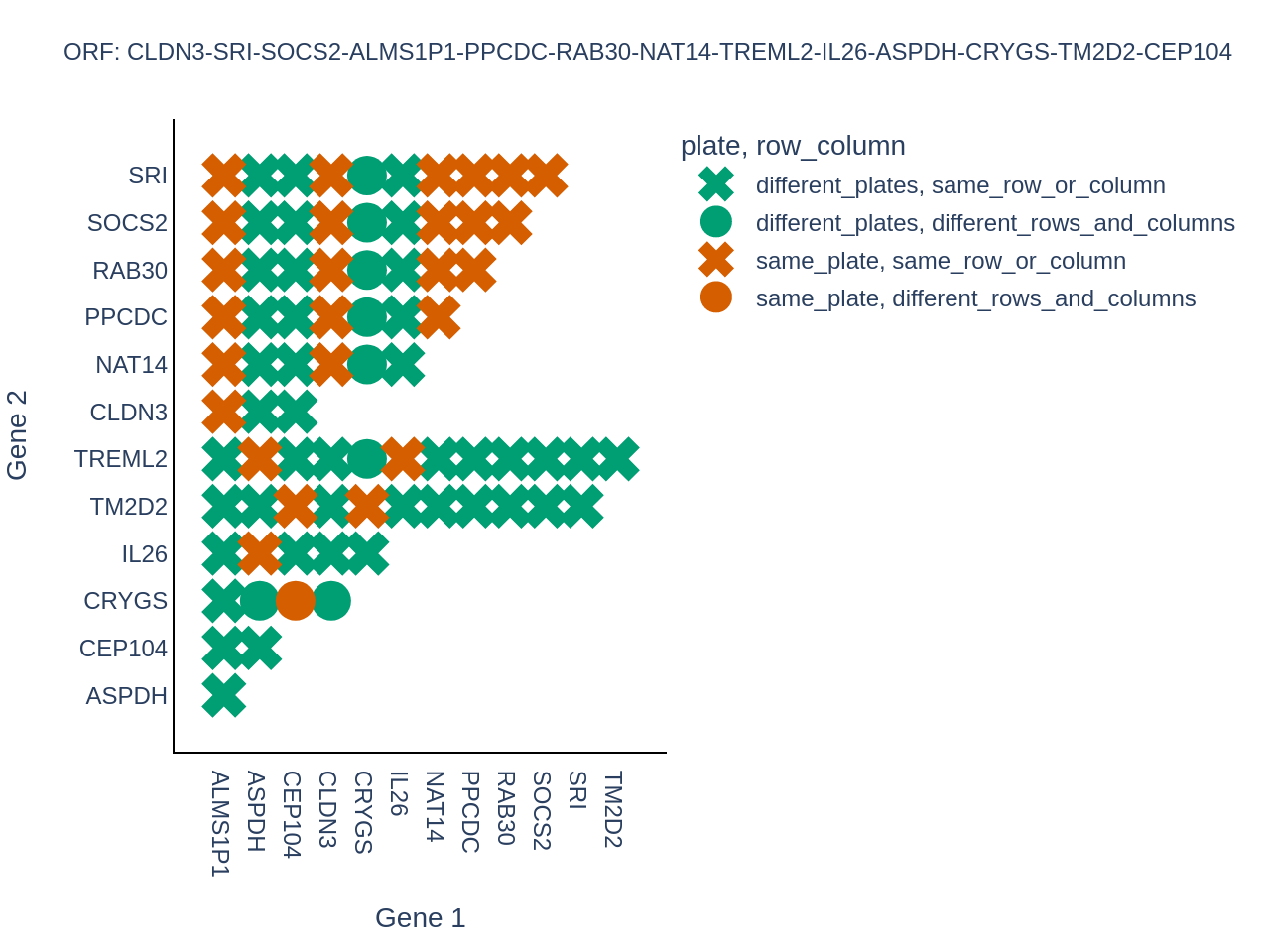

*Supplementary Figure 6**:* ***Relative location of genes in the RAB30-NAT14 cluster.*** *Most genes in this cluster are in the same row/column on the same plate or on different plates. This is an example of spurious correlation caused by plate layout effects (treatments in the same row/column on the same plate or across plates have similar phenotypes, irrespective of the identity of the treatments).*

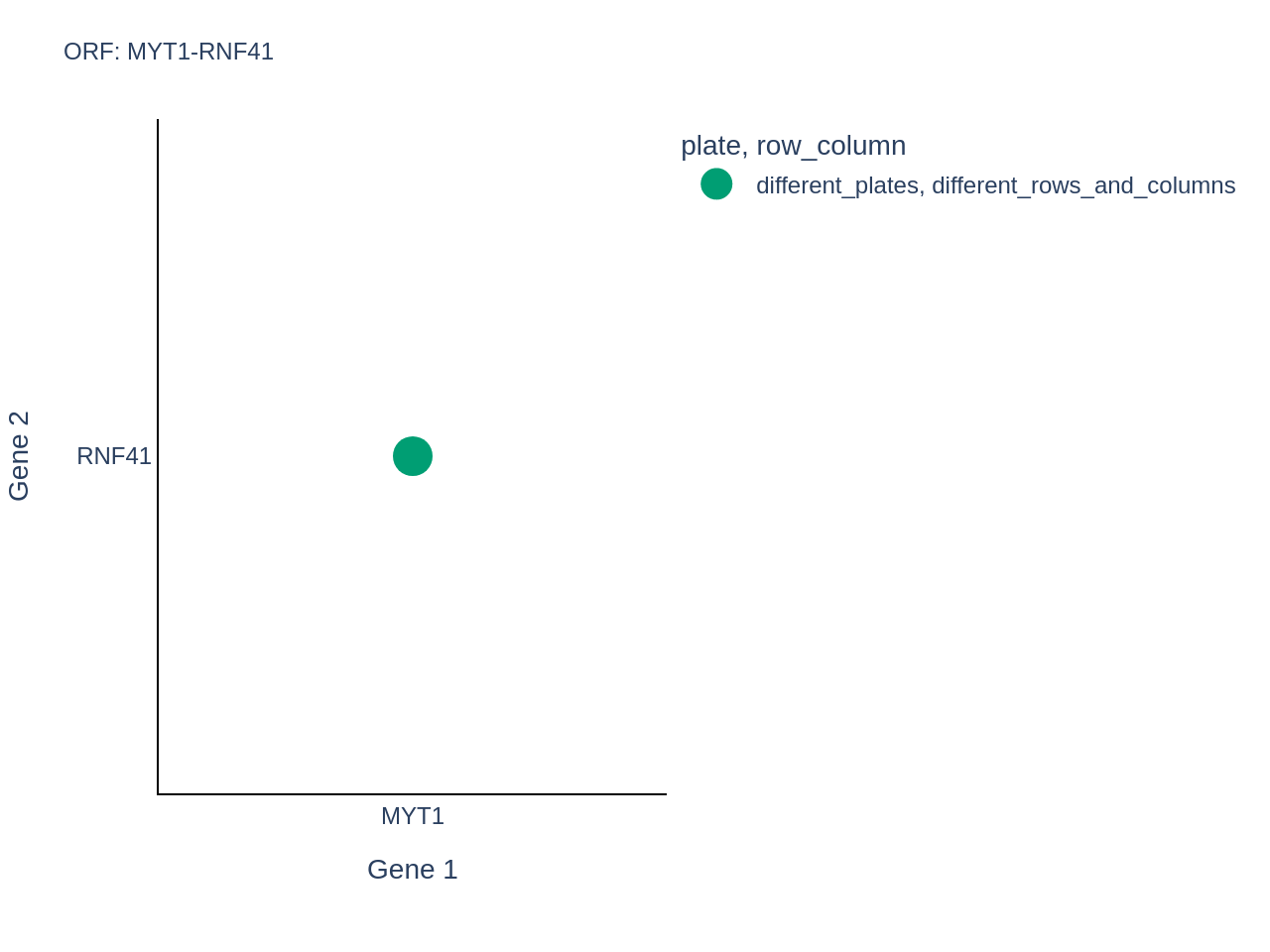

*Supplementary Figure 7**:* ***Relative location of genes in the MYT1-RNF41 cluster in the ORF dataset.***

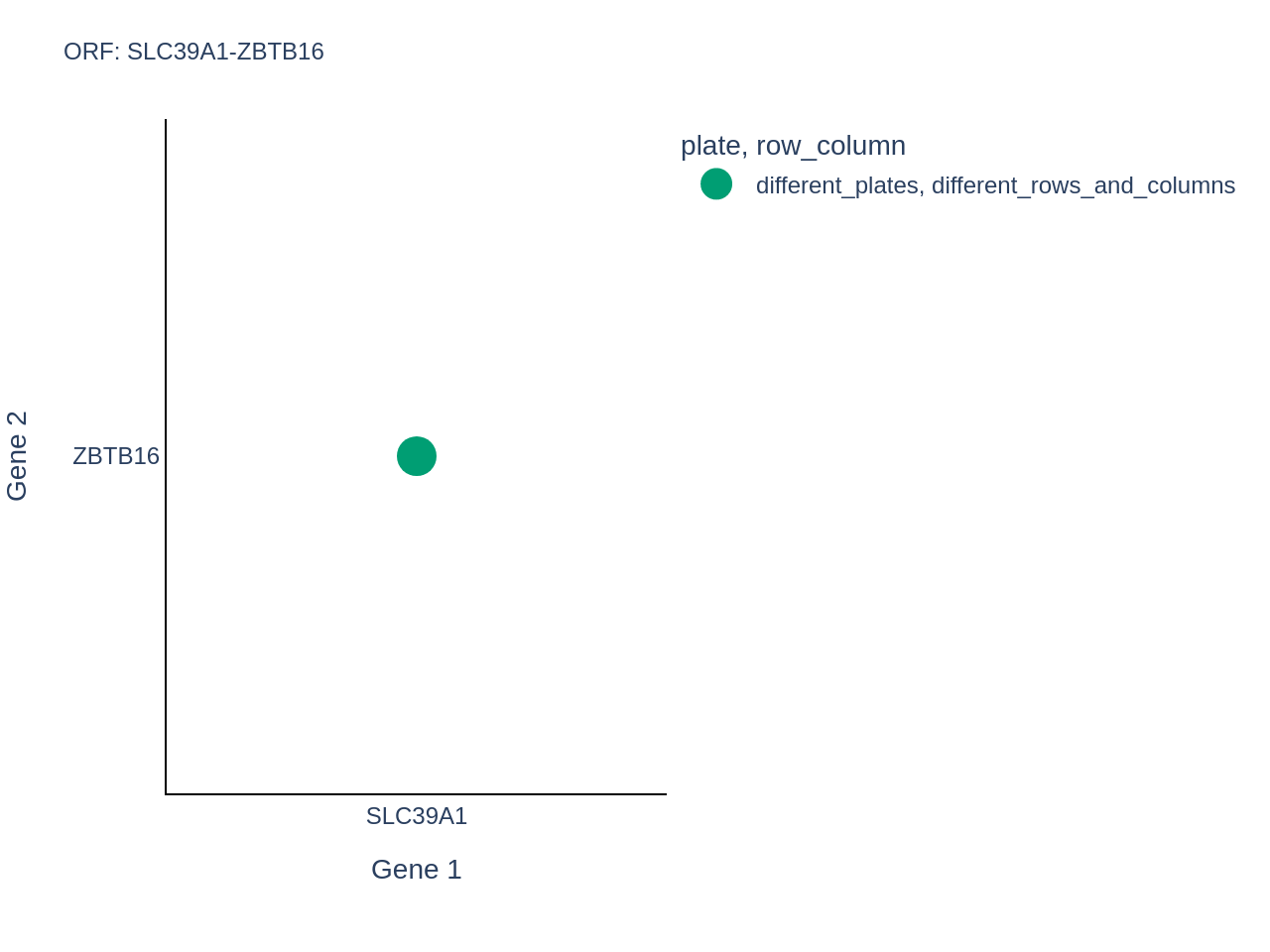

*Supplementary Figure 8**:* ***Relative location of genes in the SLC39A1-ZBTB16 cluster in the ORF dataset.***

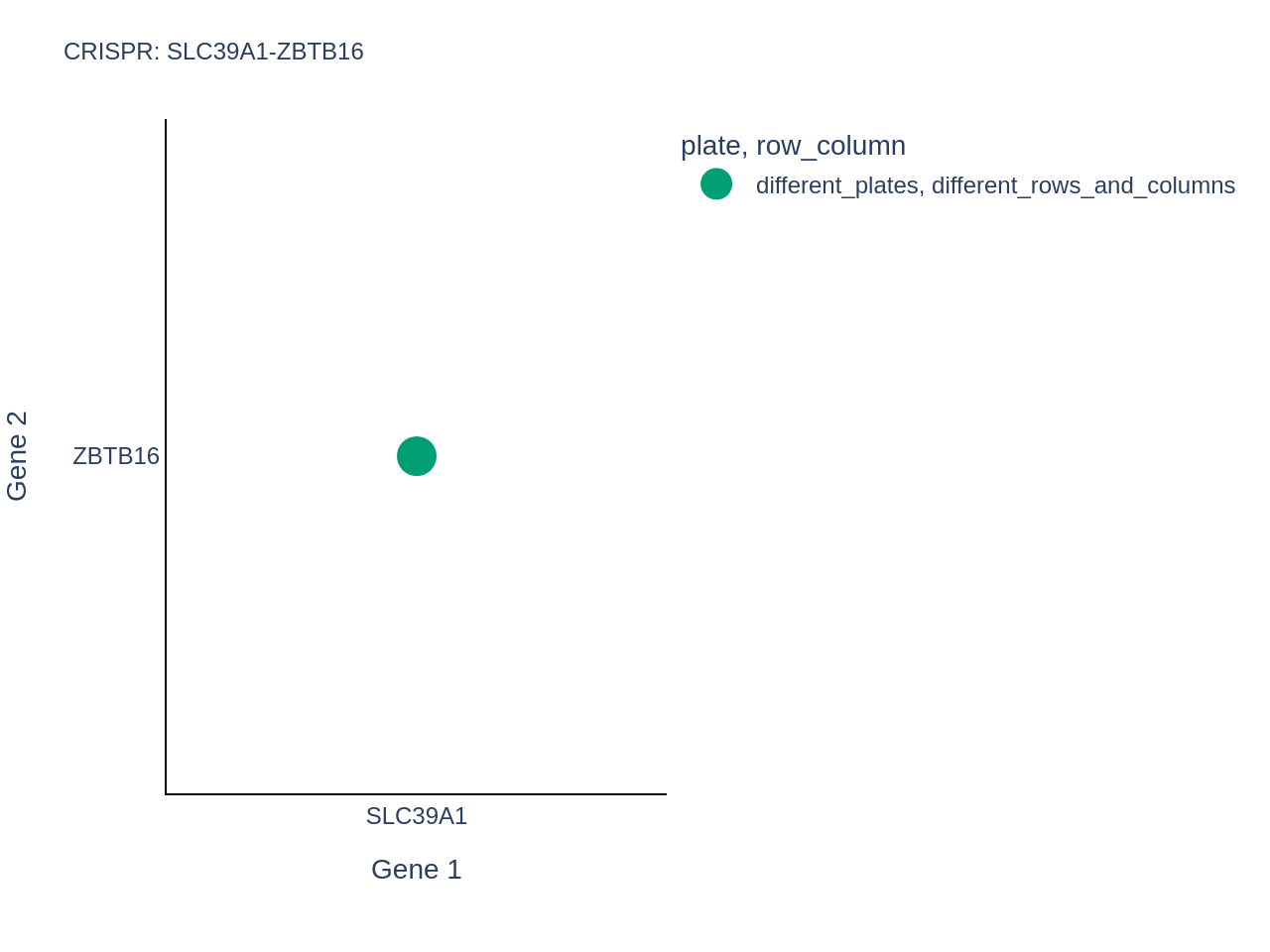

*Supplementary Figure 9**:* ***Relative location of genes in the SLC39A1-ZBTB16 cluster in the CRISPR dataset.***

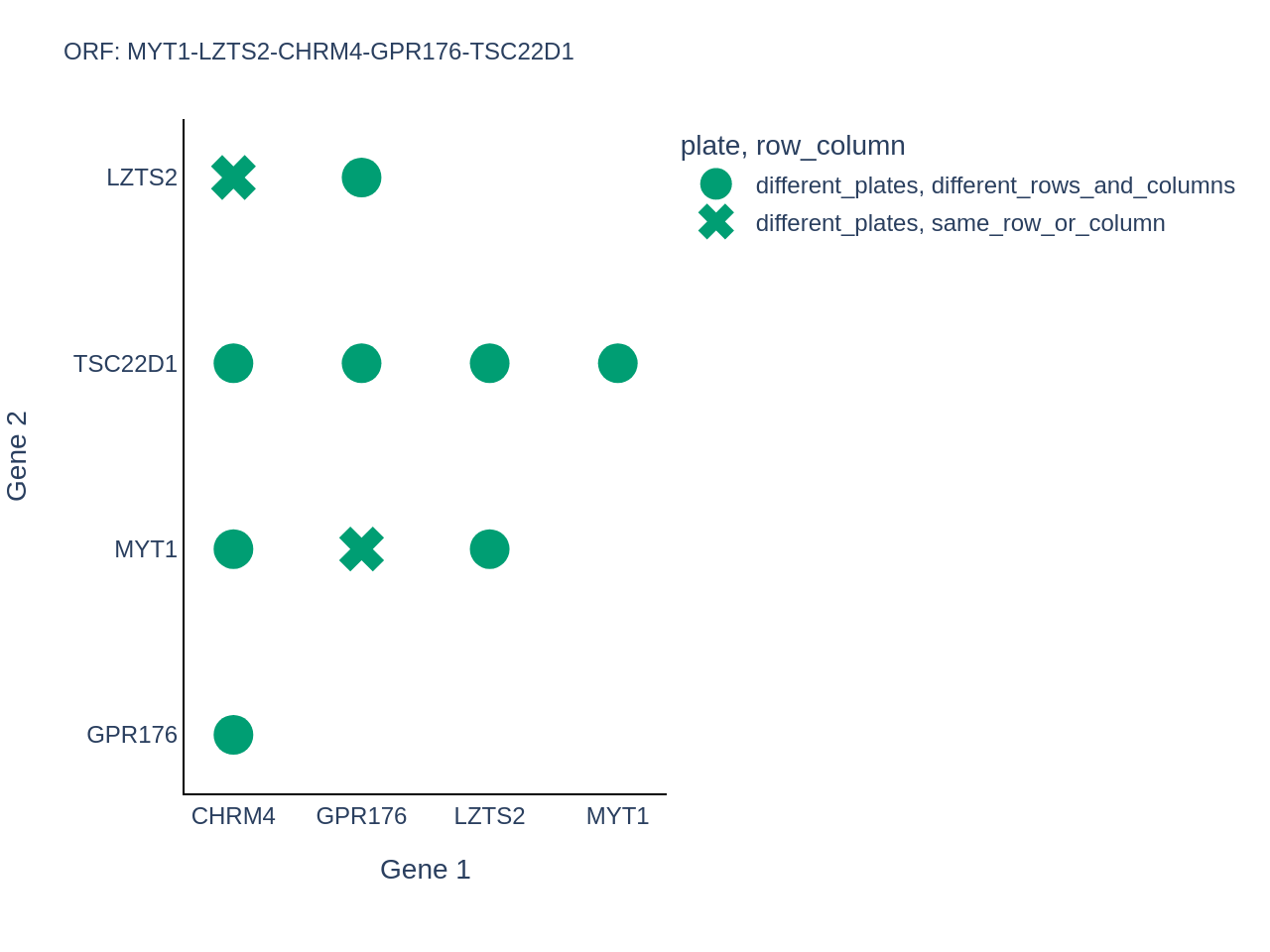

*Supplementary Figure 10**:* ***Relative location of genes in the MYT1-LZTS2-CHRM4-GPR176-TSC22D1 cluster in the ORF dataset.***

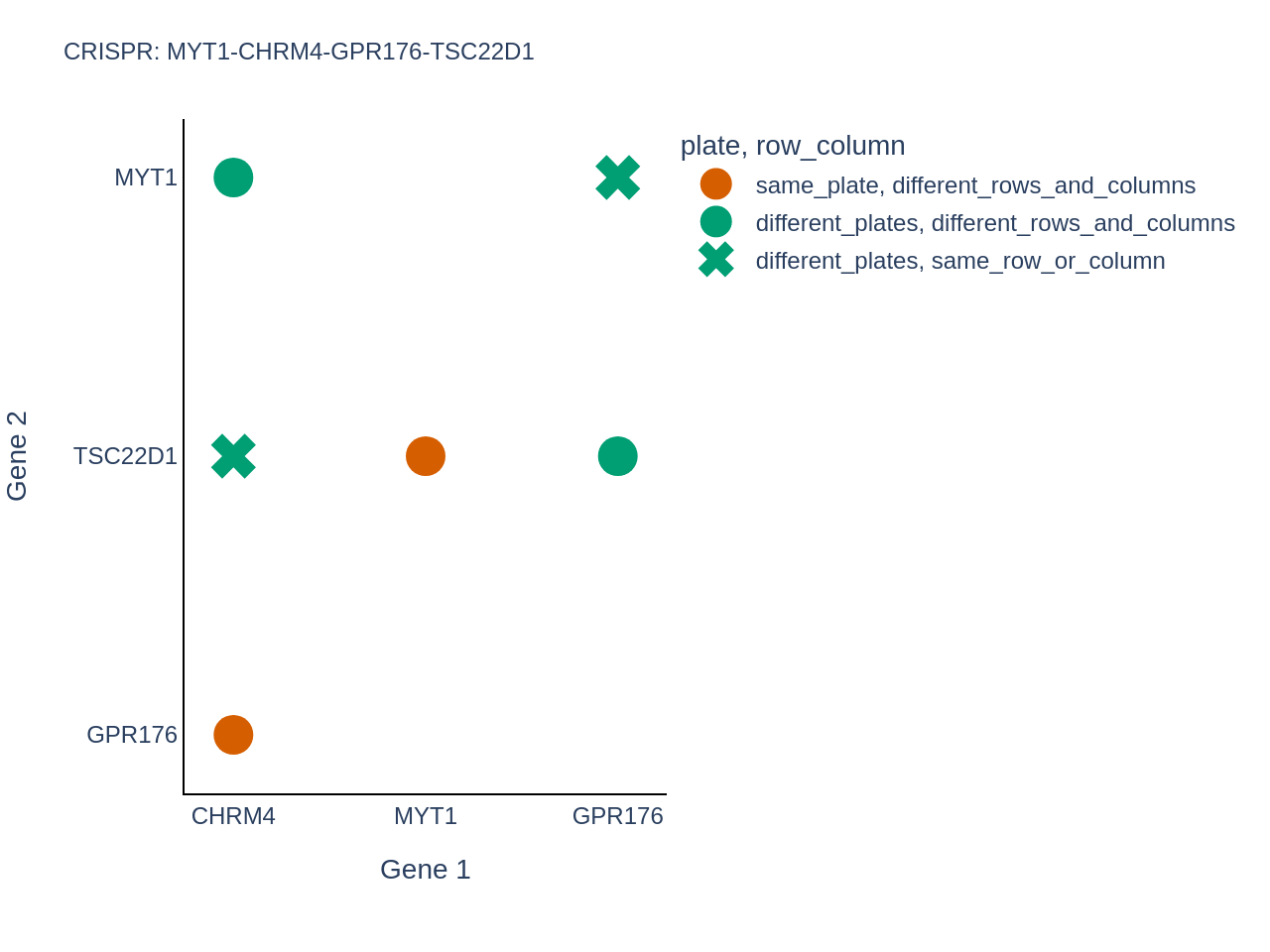

*Supplementary Figure 11**:* ***Relative location of genes in the MYT1-CHRM4-GPR176-TSC22D1 cluster in the CRISPR dataset.***

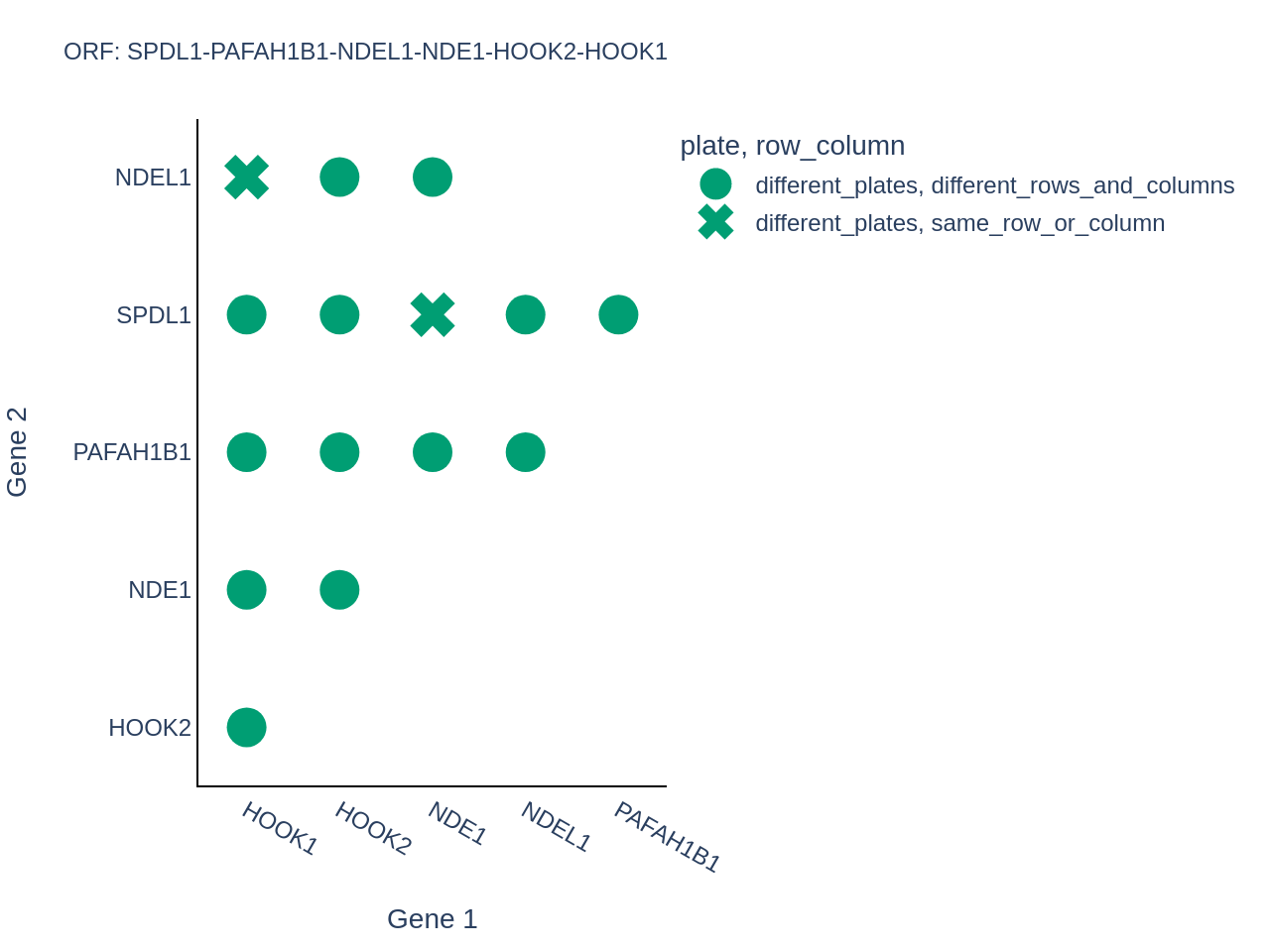

*Supplementary Figure 12**:* ***Relative location of genes in the dynein family cluster in the ORF dataset.***

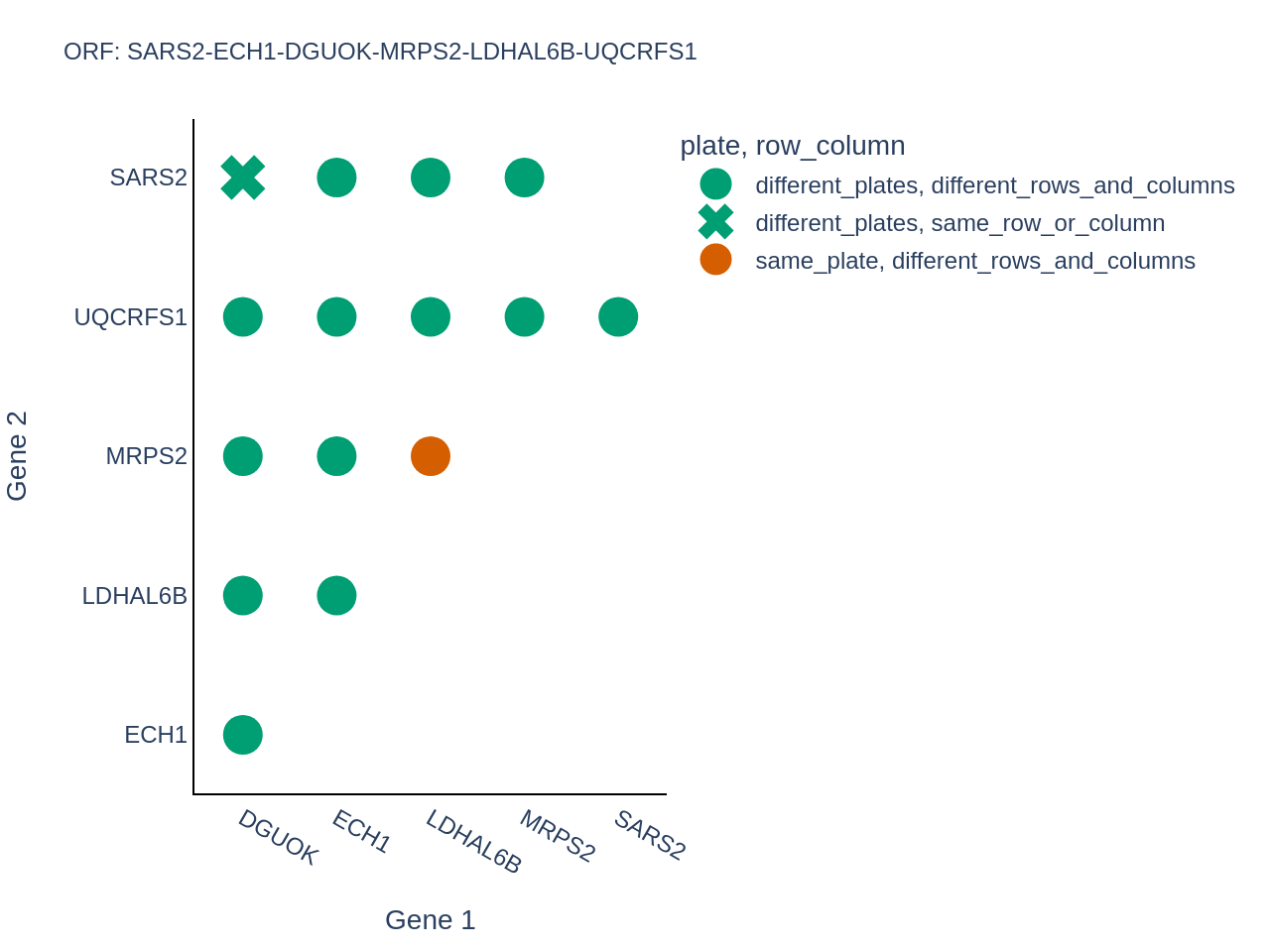

*Supplementary Figure 13**:* ***Relative location of genes in the SARS2-ECH1-DGUOK-MRSPS2-LDHAL68-UQCRFS1 cluster in the ORF dataset.***

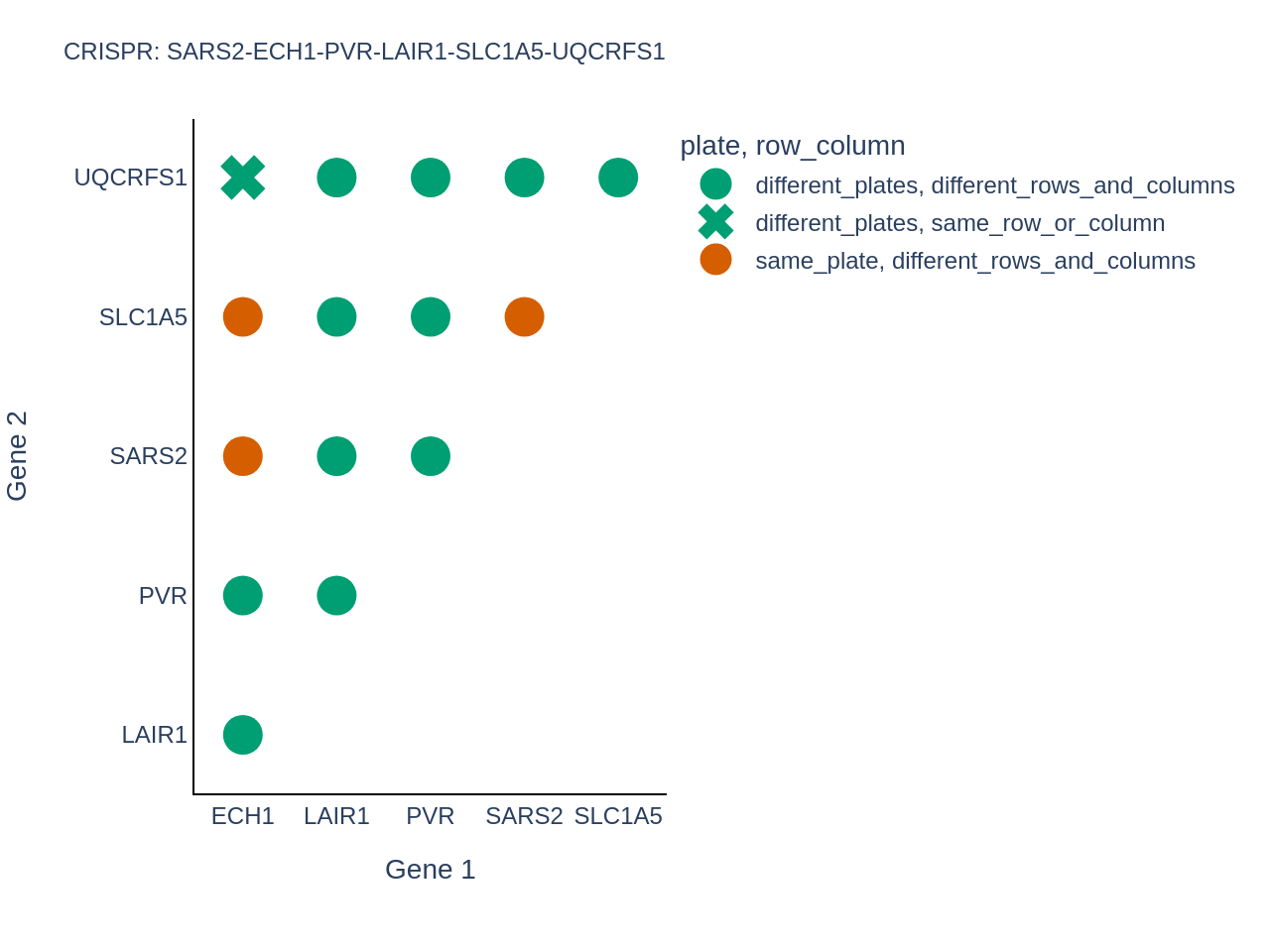

*Supplementary Figure 14**:* ***Relative location of genes in the SARS2-ECH1-PVR-LAIR1-SLC1A5-UQCRFS1 cluster in the CRISPR dataset.***

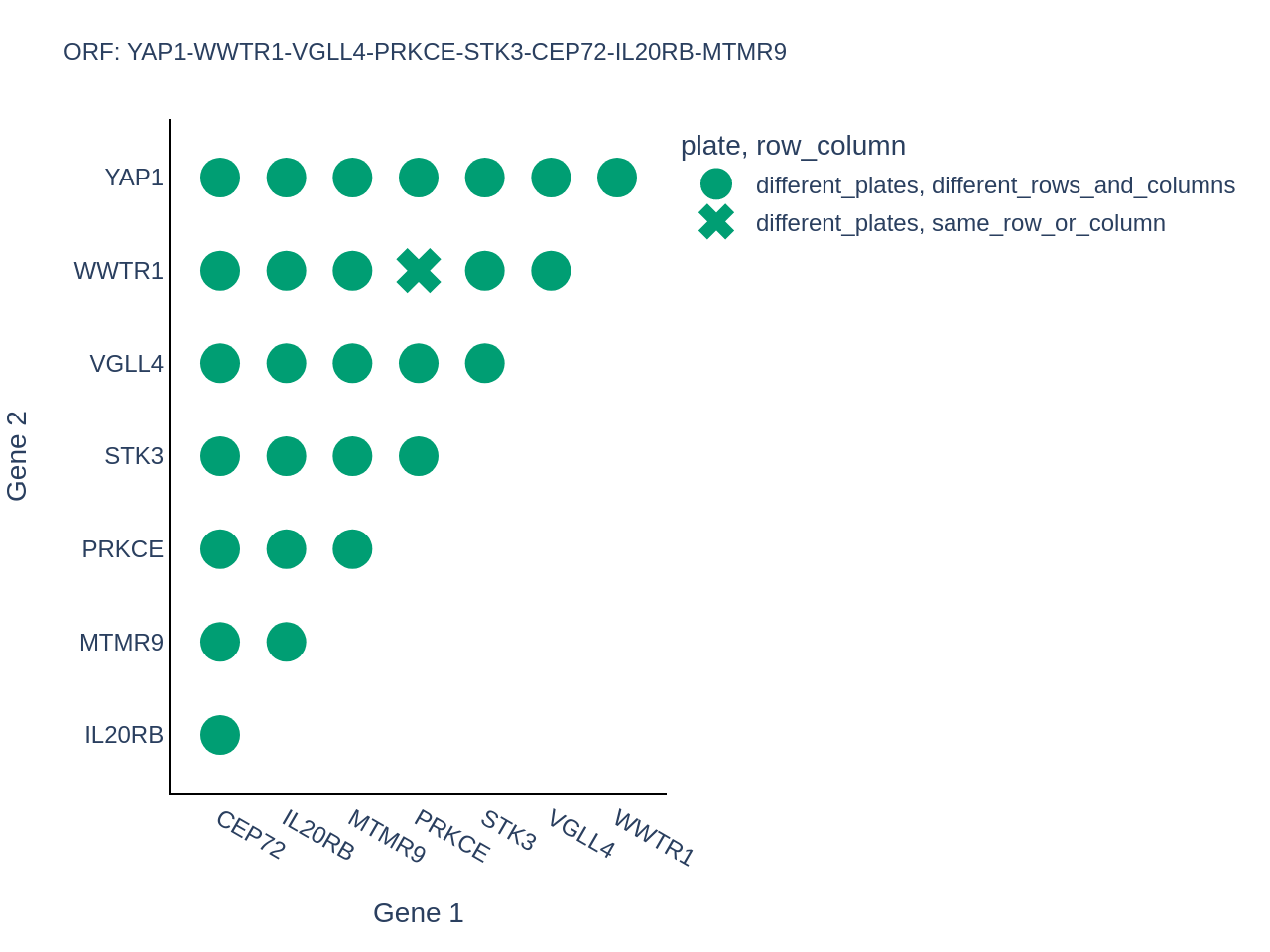

*Supplementary Figure 15**:* ***Relative location of genes in the YAP1 cluster in the ORF dataset***

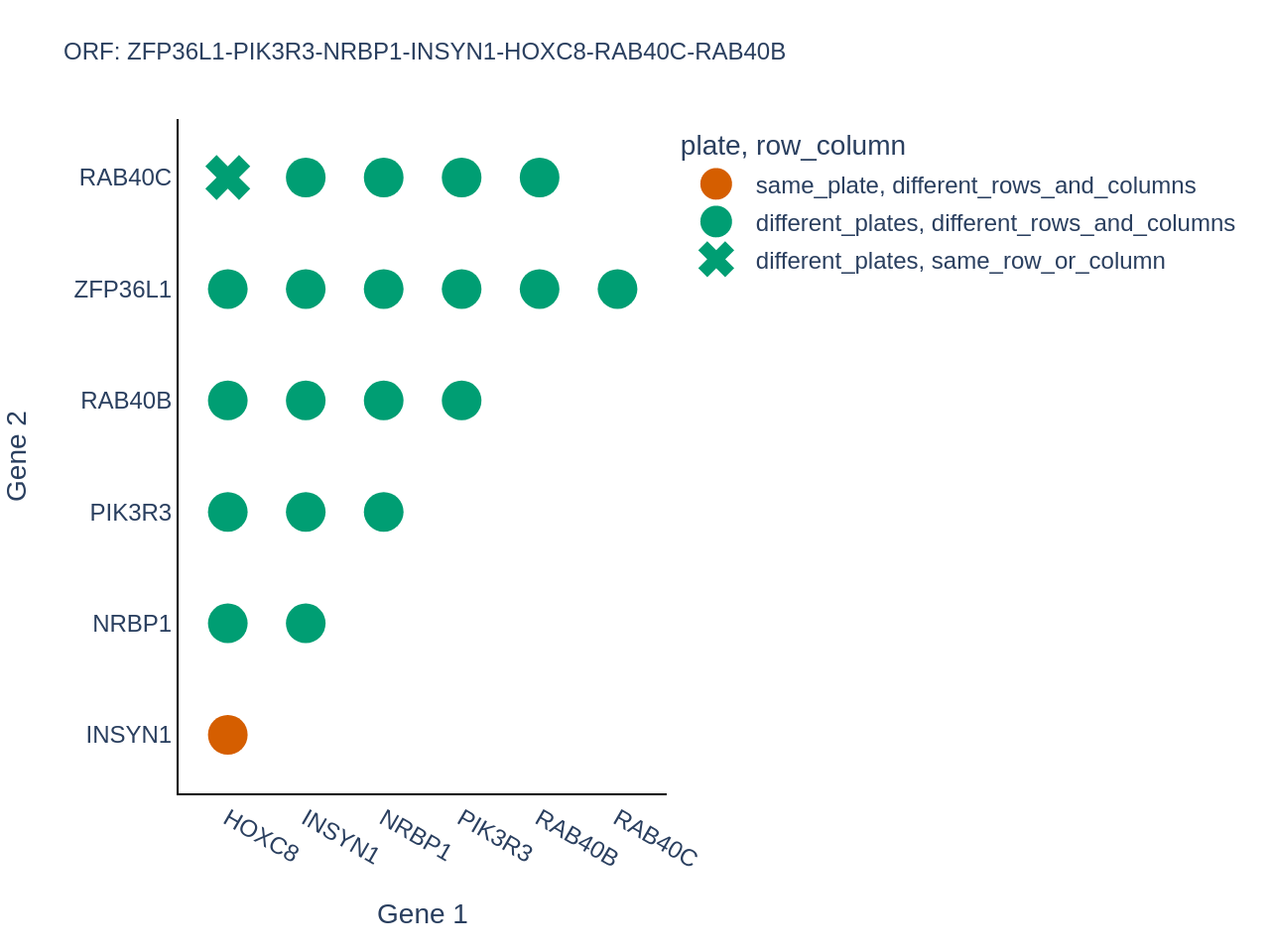

*Supplementary Figure 16**:* ***Relative location of genes in the ZFP36L1-PIK3R3-NRBP1-INSYN1-HOXC8-RAB40C-RAB40B cluster in the ORF dataset.***

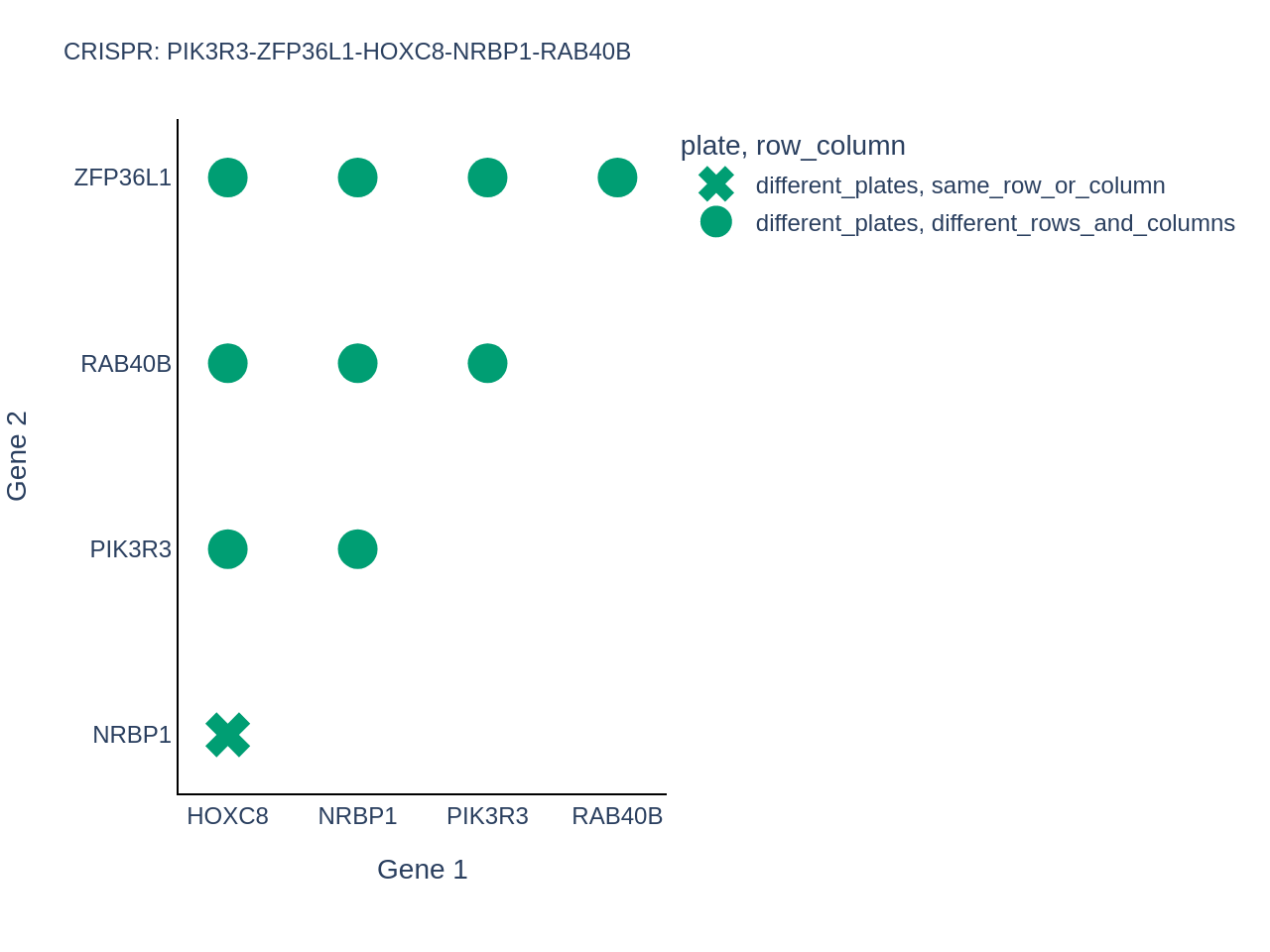

*Supplementary Figure 17**:* ***Relative location of genes in the PIK3R3-ZFP36L1-HOXC8-NRBP1-RAB40B cluster in the CRISPR dataset.***

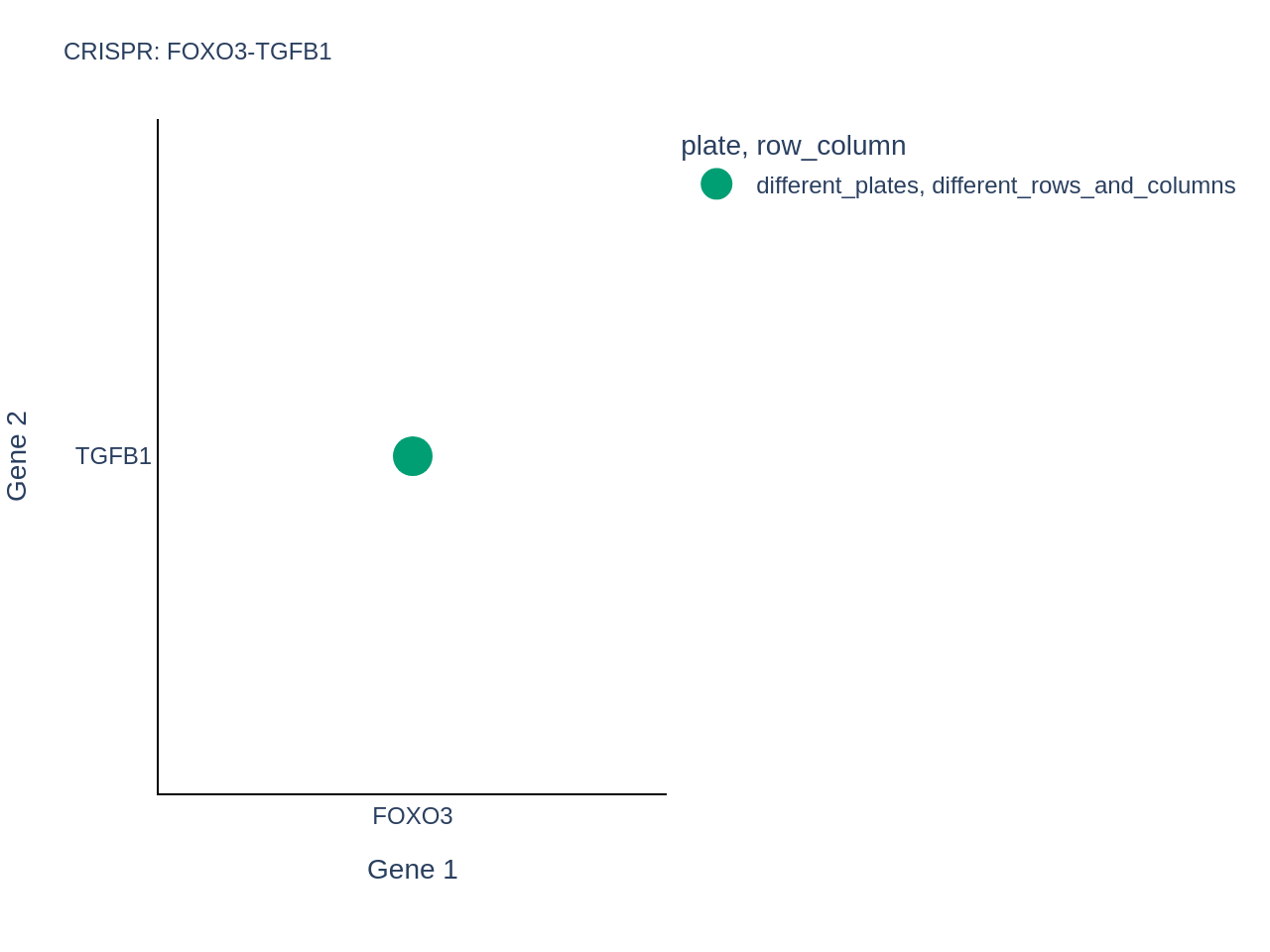

*Supplementary Figure 18**:* ***Relative location of genes in the FOXO3-TGFB1 cluster in the CRISPR dataset.***

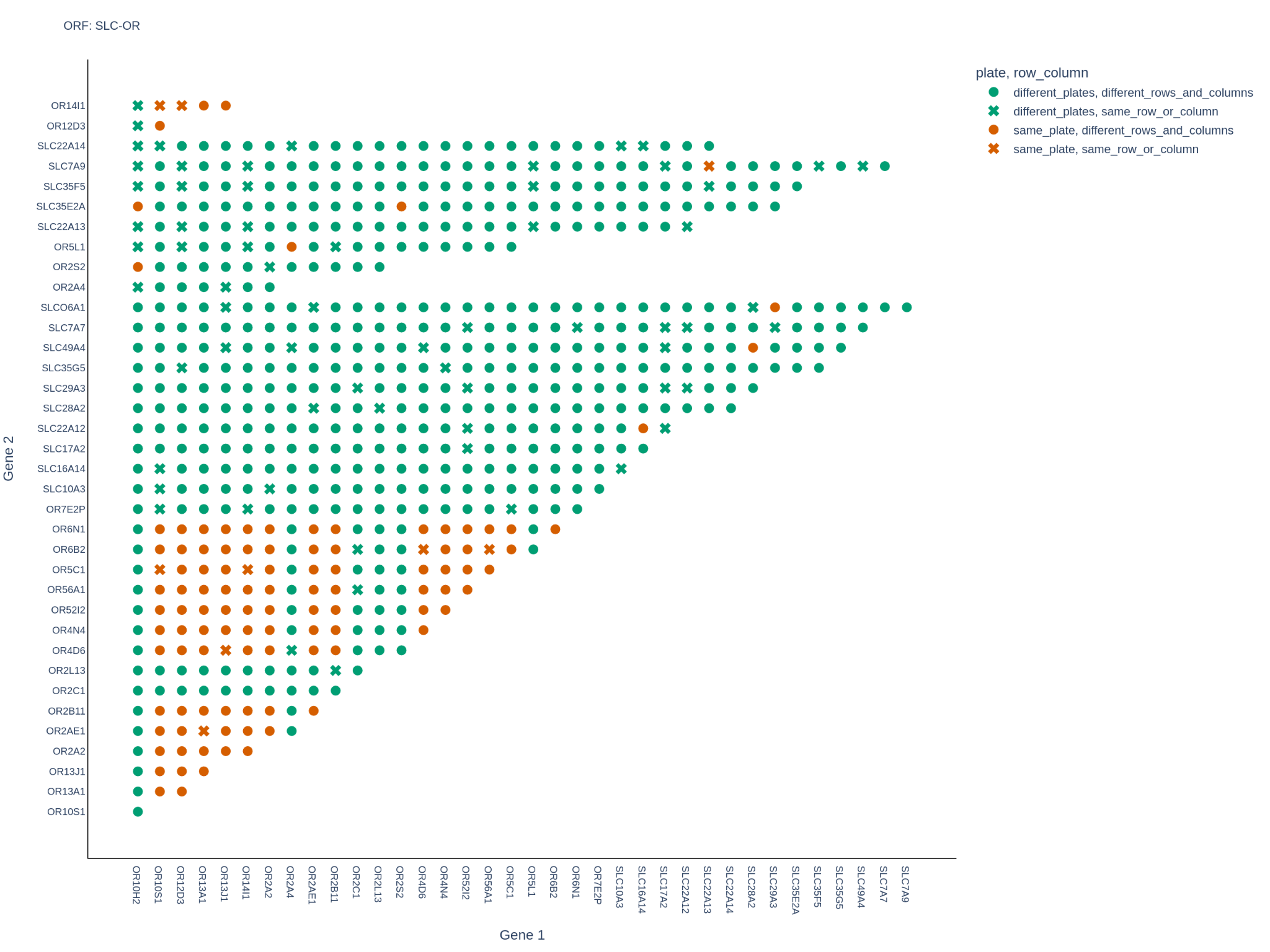

*Supplementary Figure 19**:* ***Relative location of genes in the SLC-OR cluster in the ORF dataset.*** *There aren’t any SLC-OR gene pairs that are present in the same row/column on the same plate as each other. Hence, this SLC-OR relationship cannot be attributed to spurious correlations.*

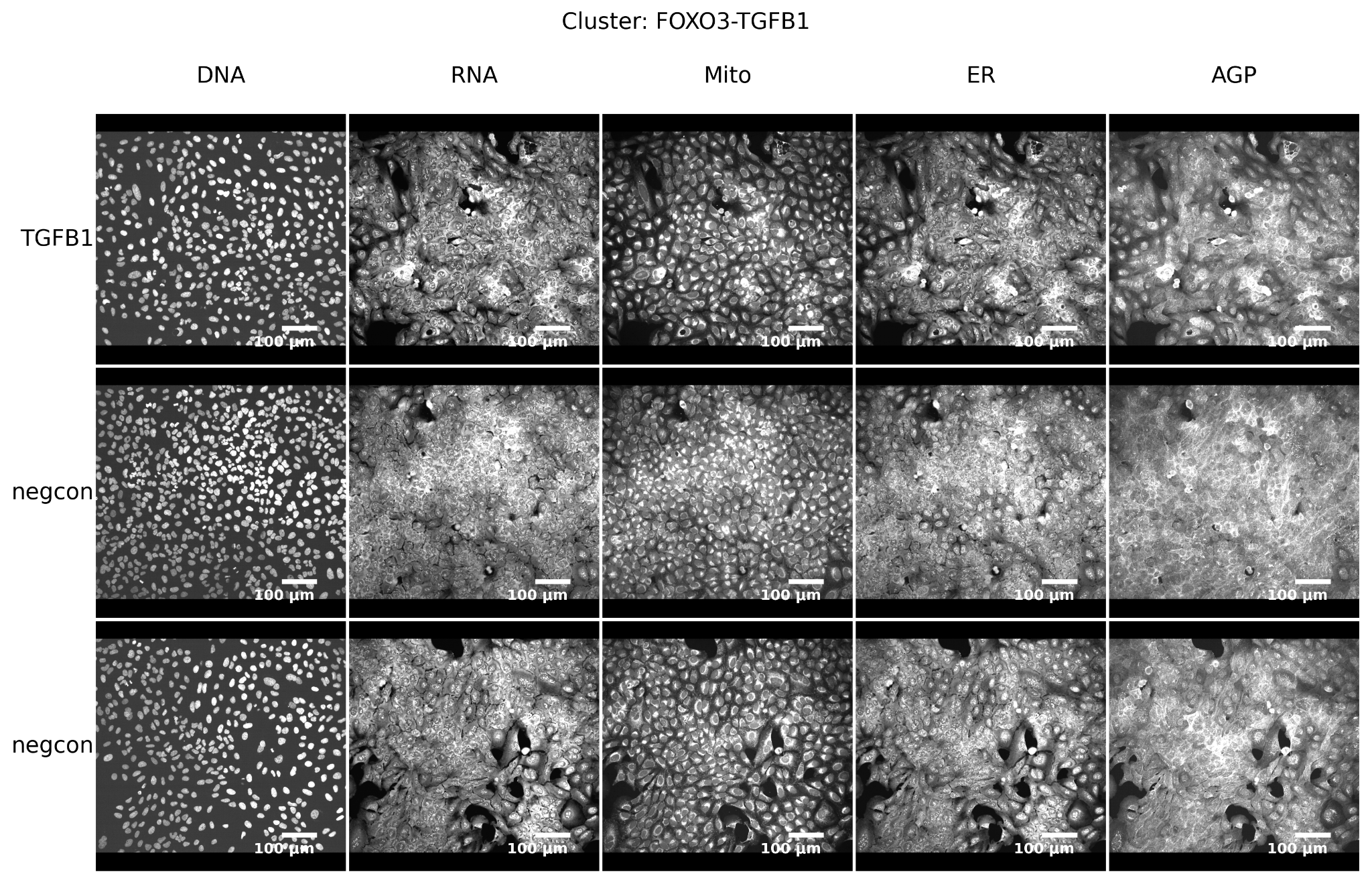

*Supplementary Figure 20**:* ***Example images from the FOXO3-TGFB1 cluster in the CRISPR dataset.*** *Representative Cell Painting images for a randomly selected gene within the cluster (top row) and a randomly selected negative control of each kind, non-targeting and no guides (bottom two rows).*

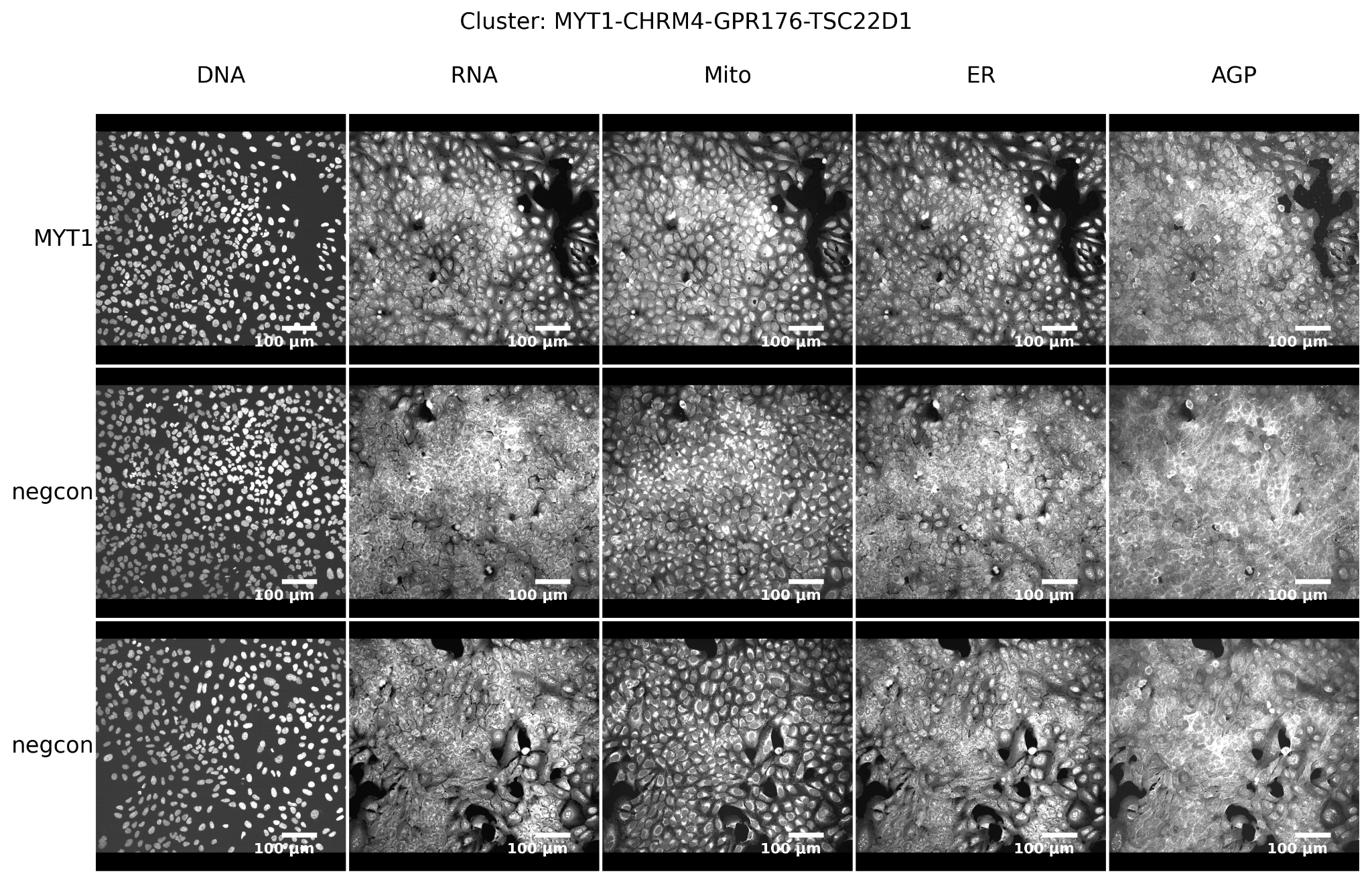

*Supplementary Figure 21**:* ***Example images from the MYT1-CHRM4-GPR176-TSC22D1 cluster in the CRISPR dataset.*** *Representative Cell Painting images for a randomly selected gene within the cluster (top row) and a randomly selected negative control of each kind, non-targeting and no guides (bottom two rows).*

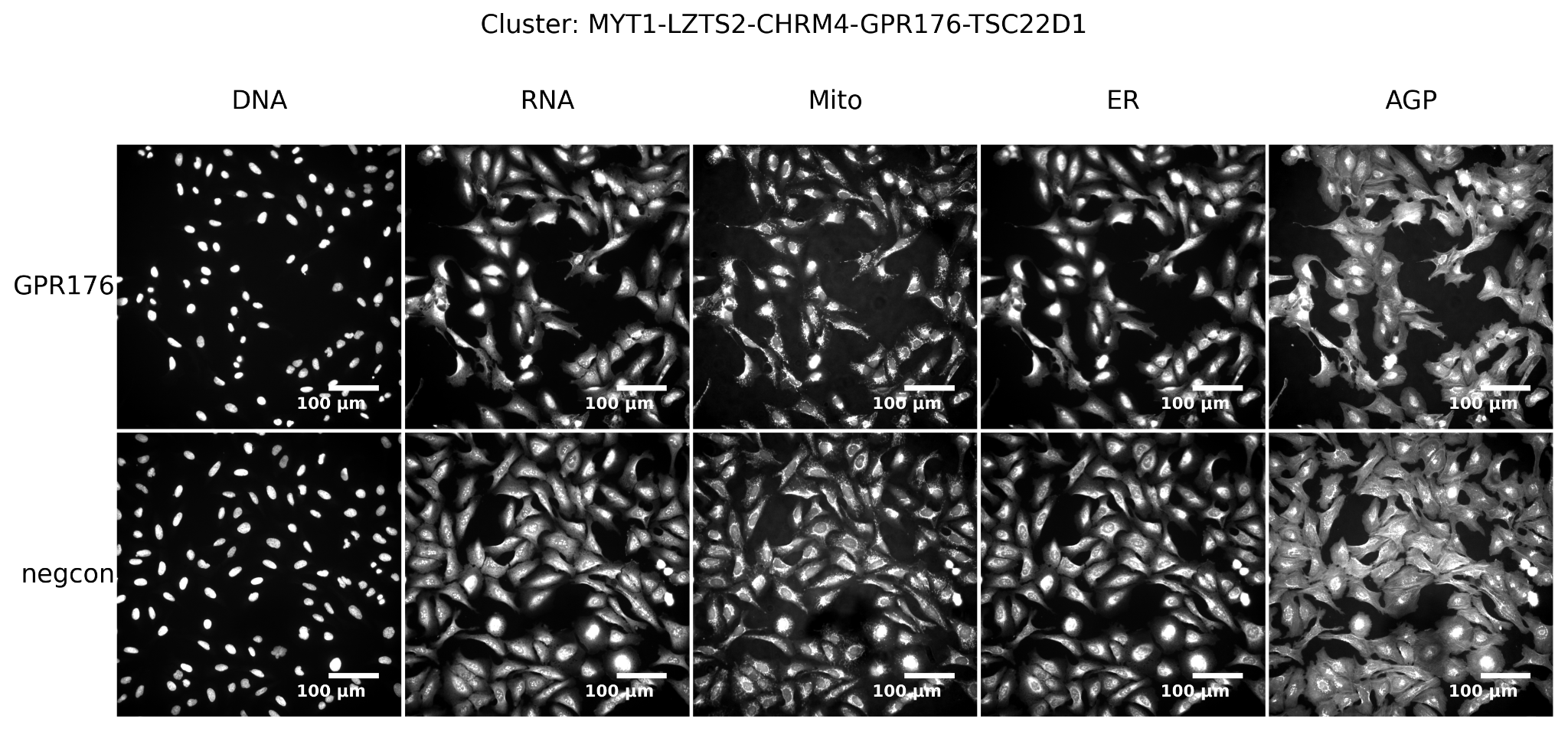

*Supplementary Figure 22**:* ***Example images from the MYT1-LZTS2-CHRM4-GPR176-TSC22D1 cluster in the ORF dataset.*** *Representative Cell Painting images for a randomly selected gene within the cluster (top row) and a randomly selected negative control (bottom row).*

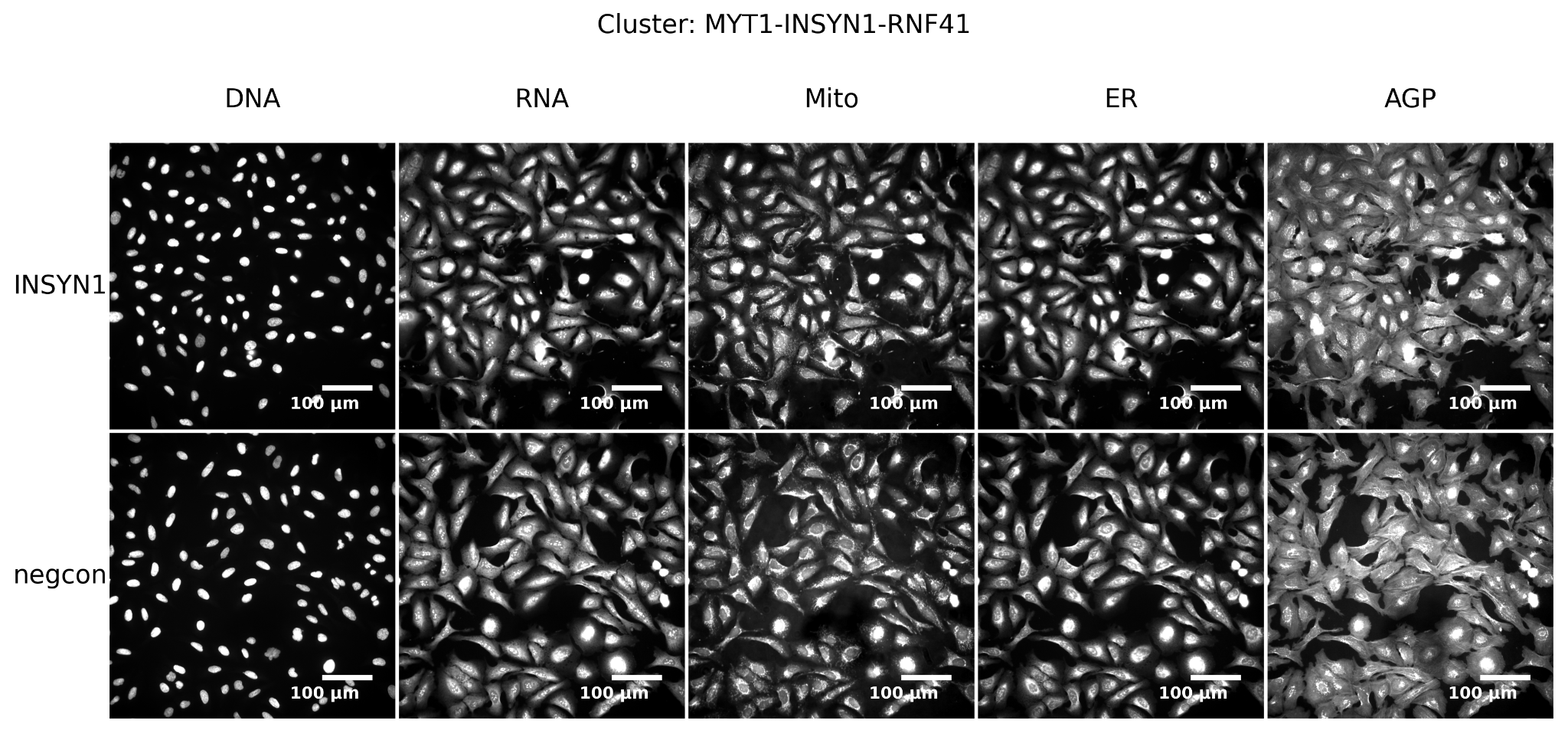

*Supplementary Figure 23**:* ***Example images from the MYT1-RNF41 cluster in the ORF dataset.*** *Representative Cell Painting images for a randomly selected gene within the cluster (top row) and a randomly selected negative control (bottom row).*

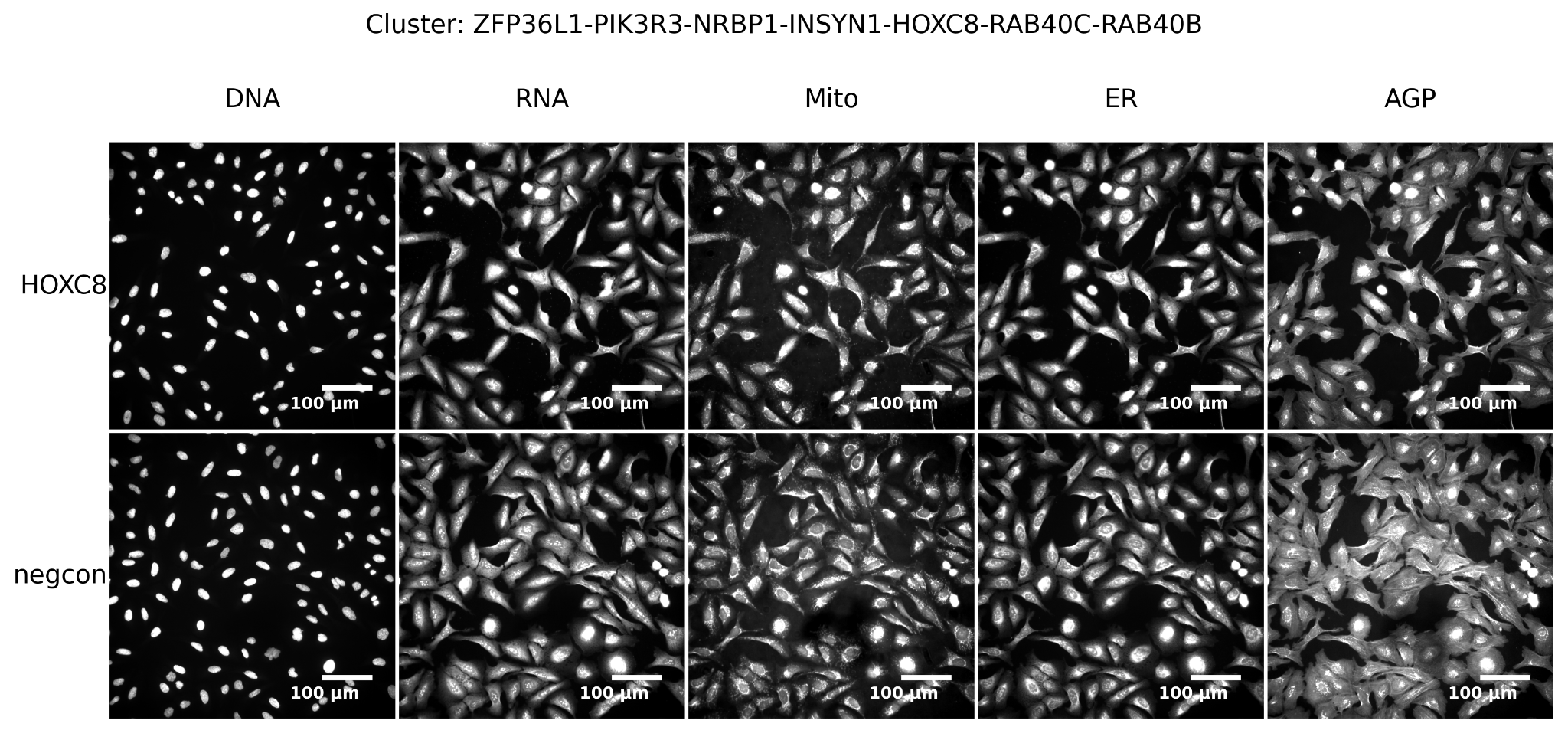

*Supplementary Figure 24**:* ***Example images from the ZFP36L1-PIK3R3-NRBP1-INSYN1-HOXC8-RAB40C-RAB40B cluster in the ORF dataset.*** *Representative Cell Painting images for a randomly selected gene within the cluster (top row) and a randomly selected negative control (bottom row).*

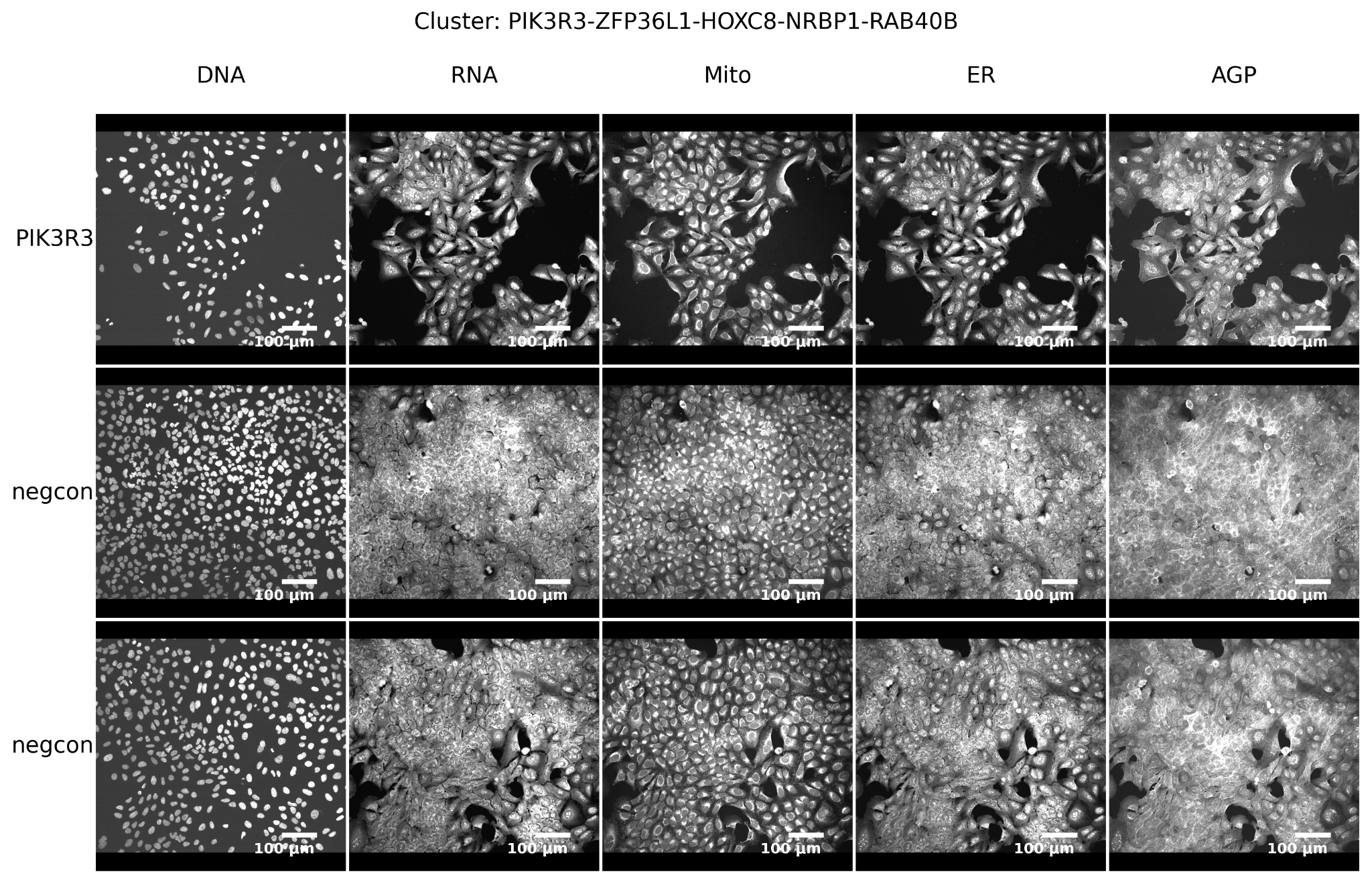

*Supplementary Figure 25**:* ***Example images from the PIK3R3-ZFP36L1-HOXC8-NRBP1-RAB40B cluster in the CRISPR dataset.*** *Representative Cell Painting images for a randomly selected gene within the cluster (top row) and a randomly selected negative control of each kind, non-targeting and no guides (bottom two rows).*

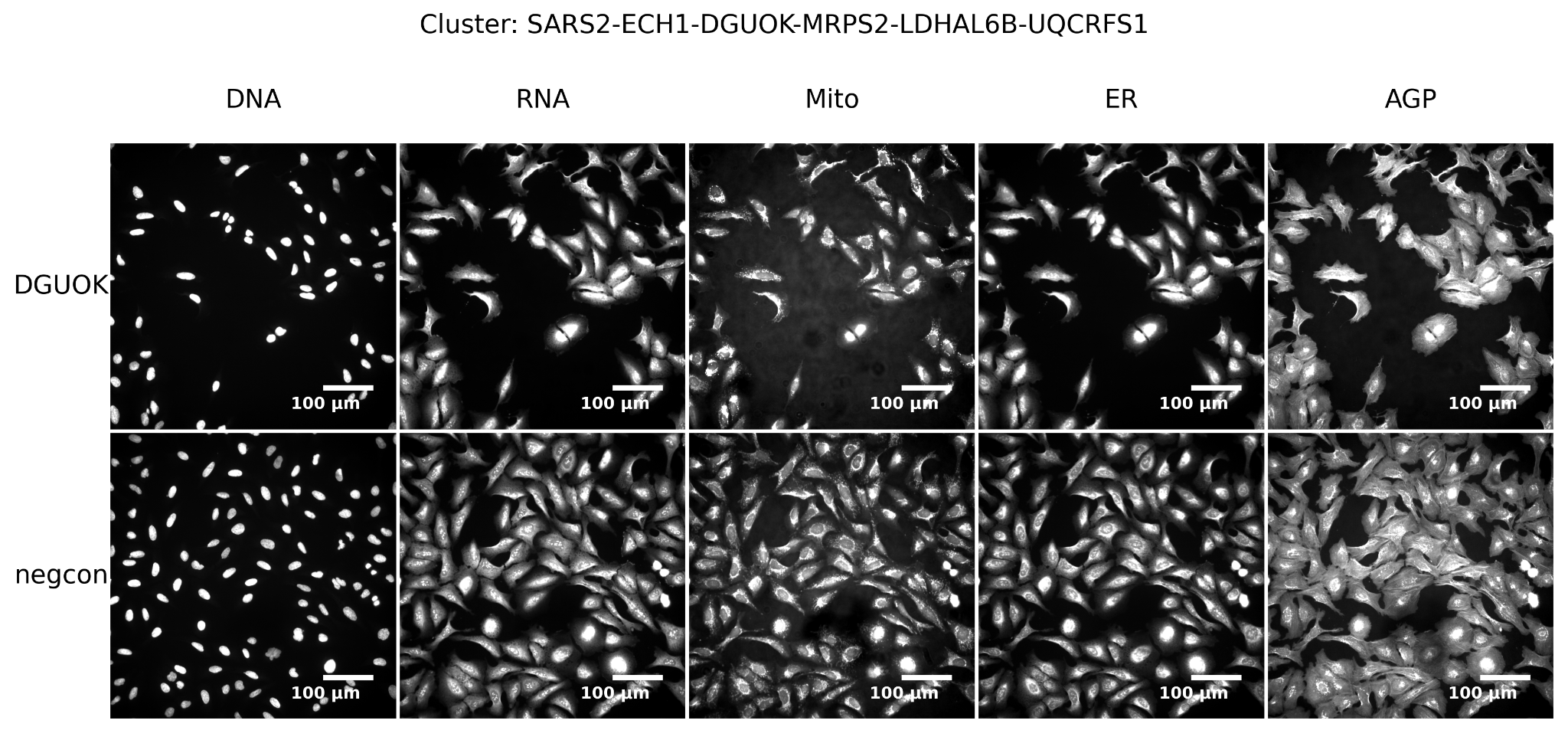

*Supplementary Figure 26**:* ***Example images from the SARS2-ECH1-DGUOK-MRSPS2-LDHAL68-UQCRFS1 cluster in the ORF dataset.*** *Representative Cell Painting images for a randomly selected gene within the cluster (top row) and a randomly selected negative control (bottom row).*

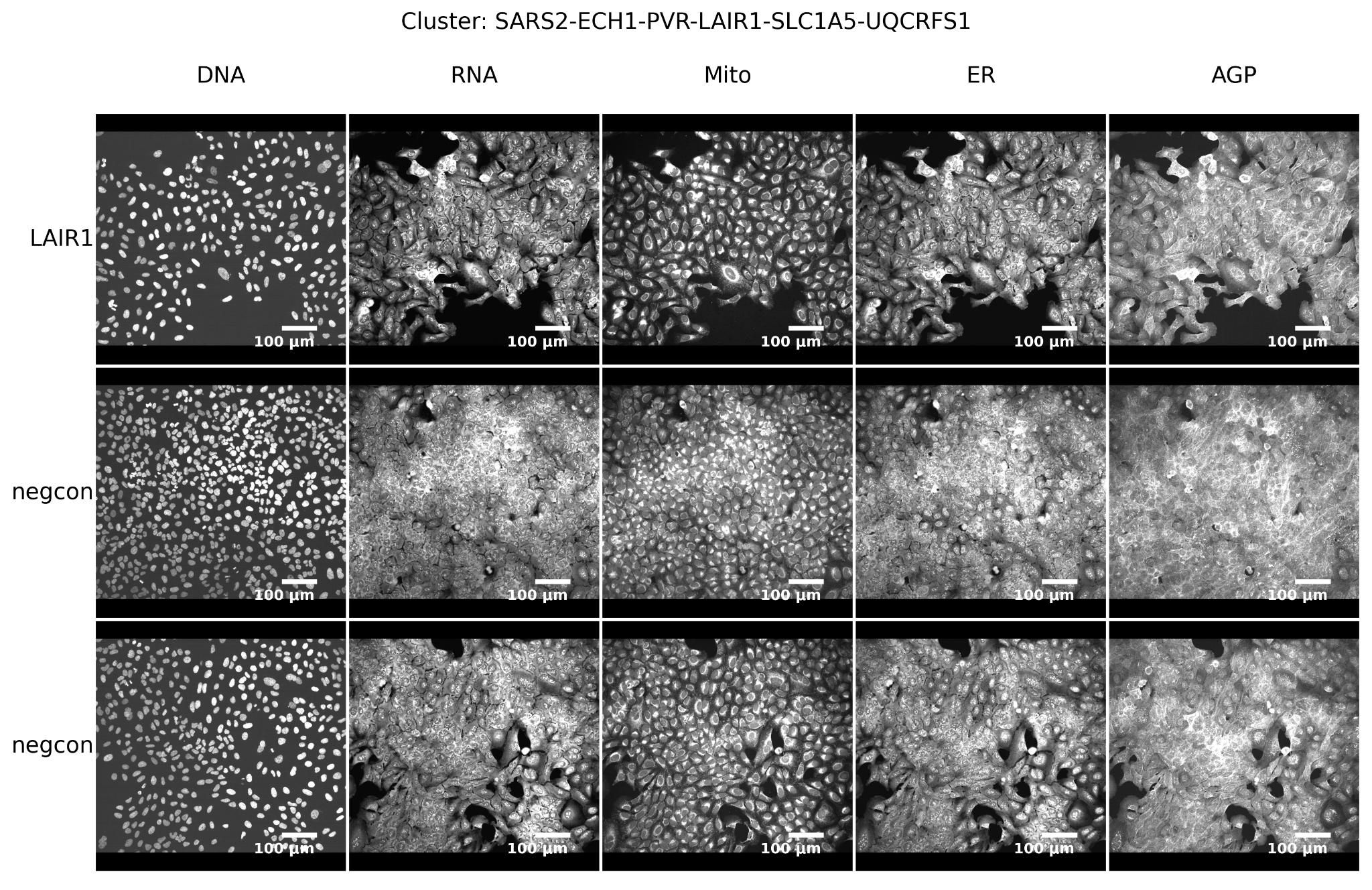

*Supplementary Figure 27**:* ***Example images from the SARS2-ECH1-PVR-LAIR1-SLC1A5-UQCRFS1 cluster in the CRISPR dataset.*** *Representative Cell Painting images for a randomly selected gene within the cluster (top row) and a randomly selected negative control of each kind, non-targeting and no guides (bottom two rows).*

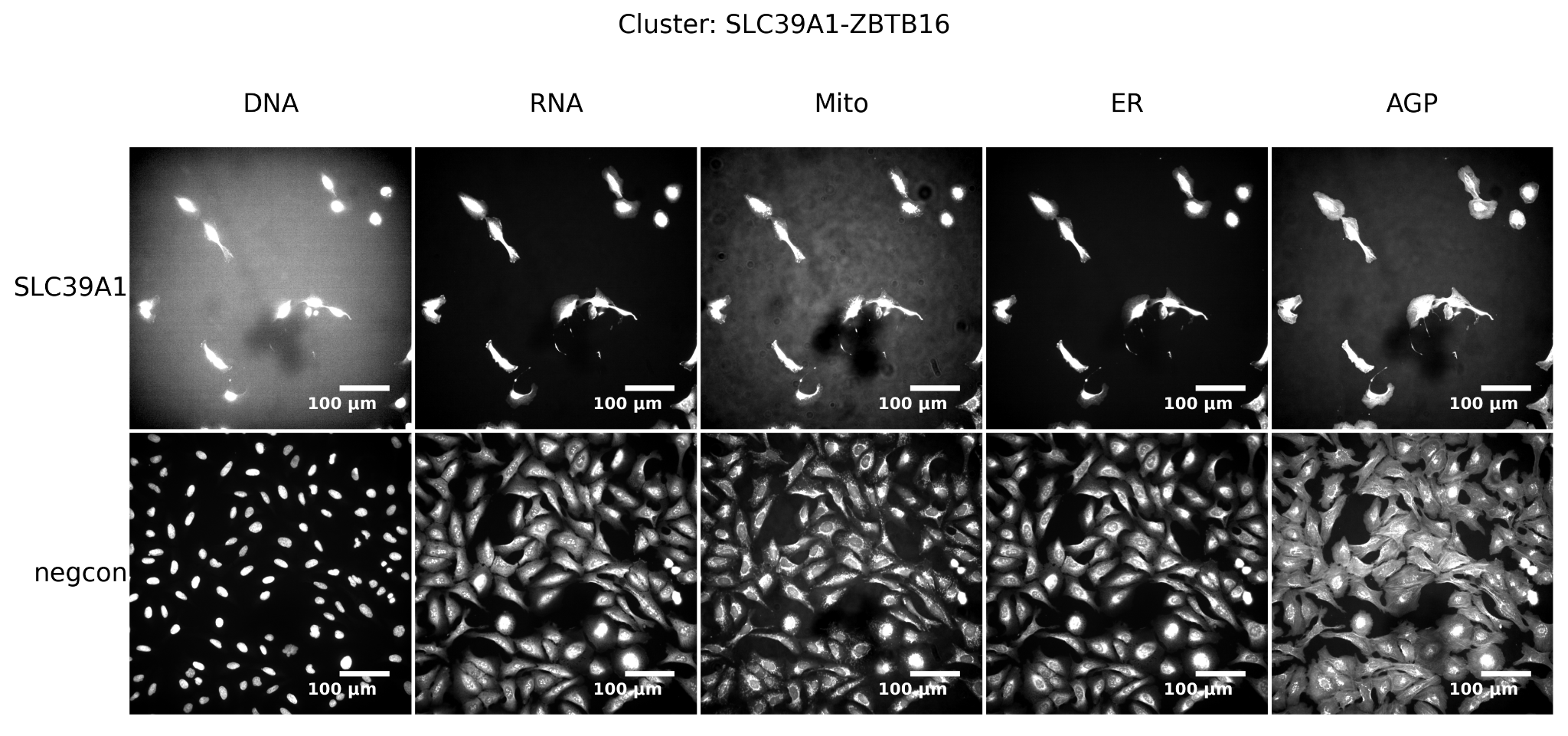

*Supplementary Figure 28**:* ***Example images from the SLC39A1-ZBTB16 cluster in the ORF dataset.*** *Representative Cell Painting images for a randomly selected gene within the cluster (top row) and a randomly selected negative control (bottom row).*

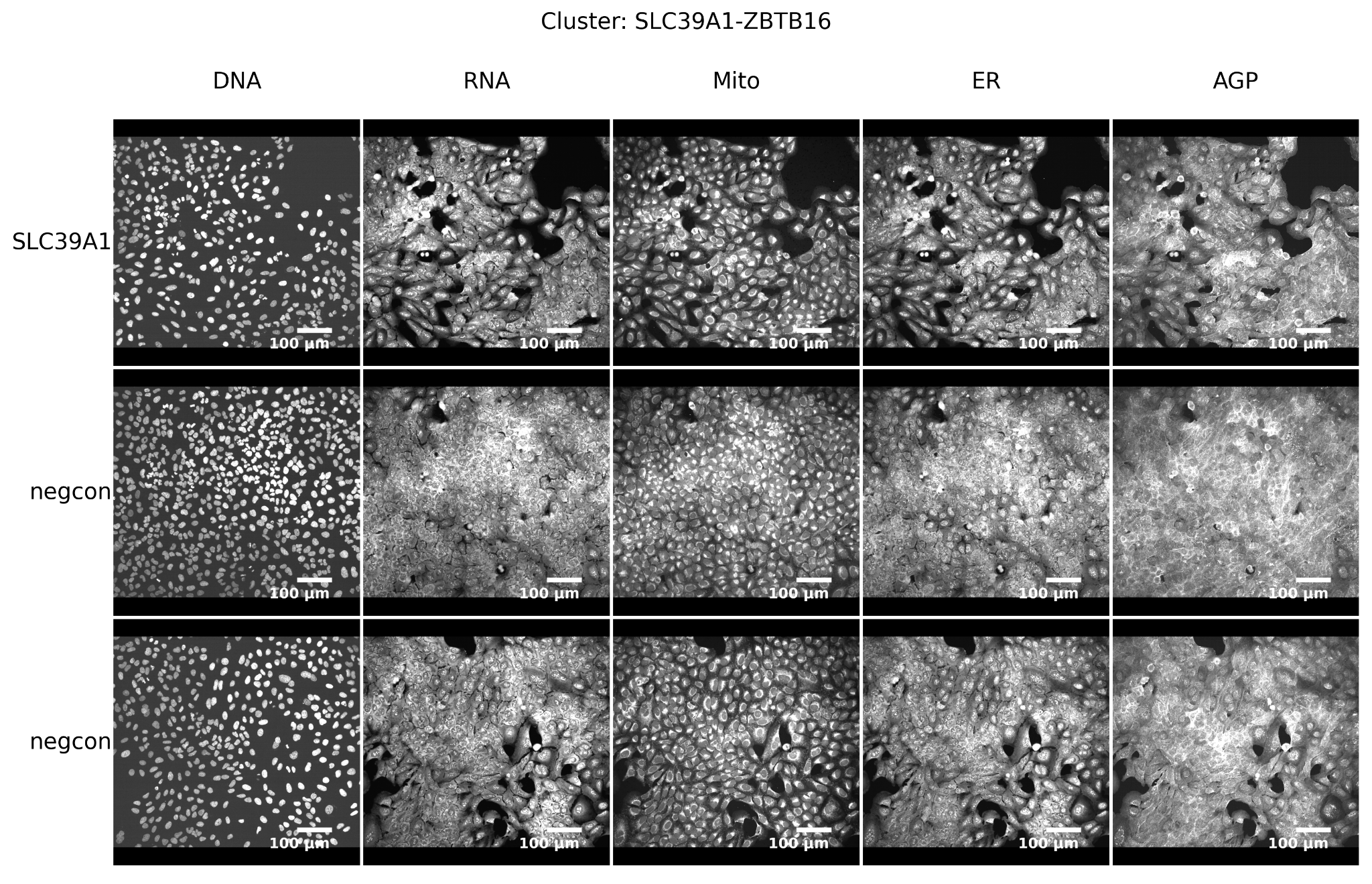

*Supplementary Figure 29**:* ***Example images from the SLC39A1-ZBTB16 cluster in the CRISPR dataset.*** *Representative Cell Painting images for a randomly selected gene within the cluster (top row) and a randomly selected negative control of each kind, non-targeting and no guides (bottom two rows).*

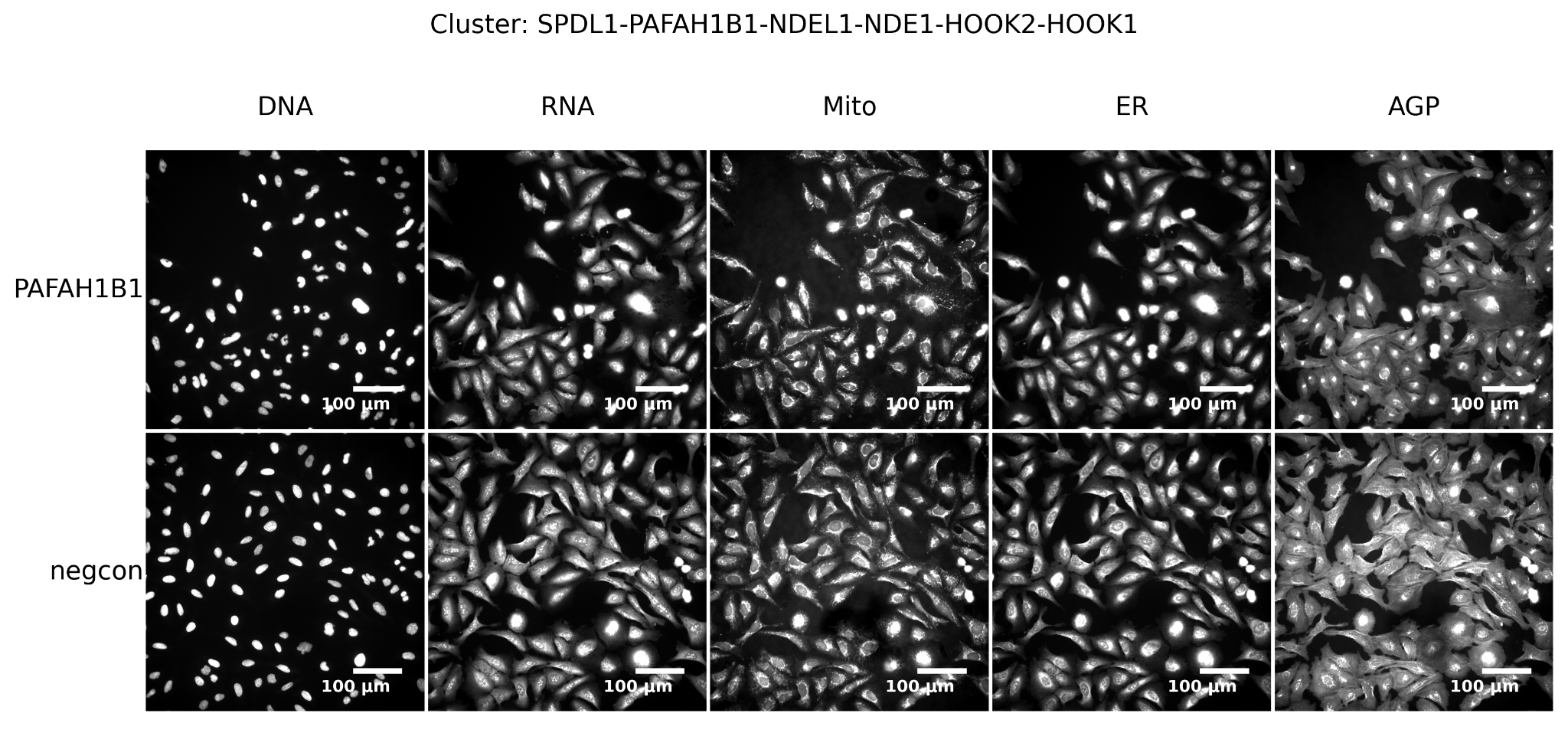
*Supplementary Figure 30**:* ***Example images from the dynein family cluster in the ORF dataset.*** *Representative Cell Painting images for a randomly selected gene within the cluster (top row) and a randomly selected negative control (bottom row).*

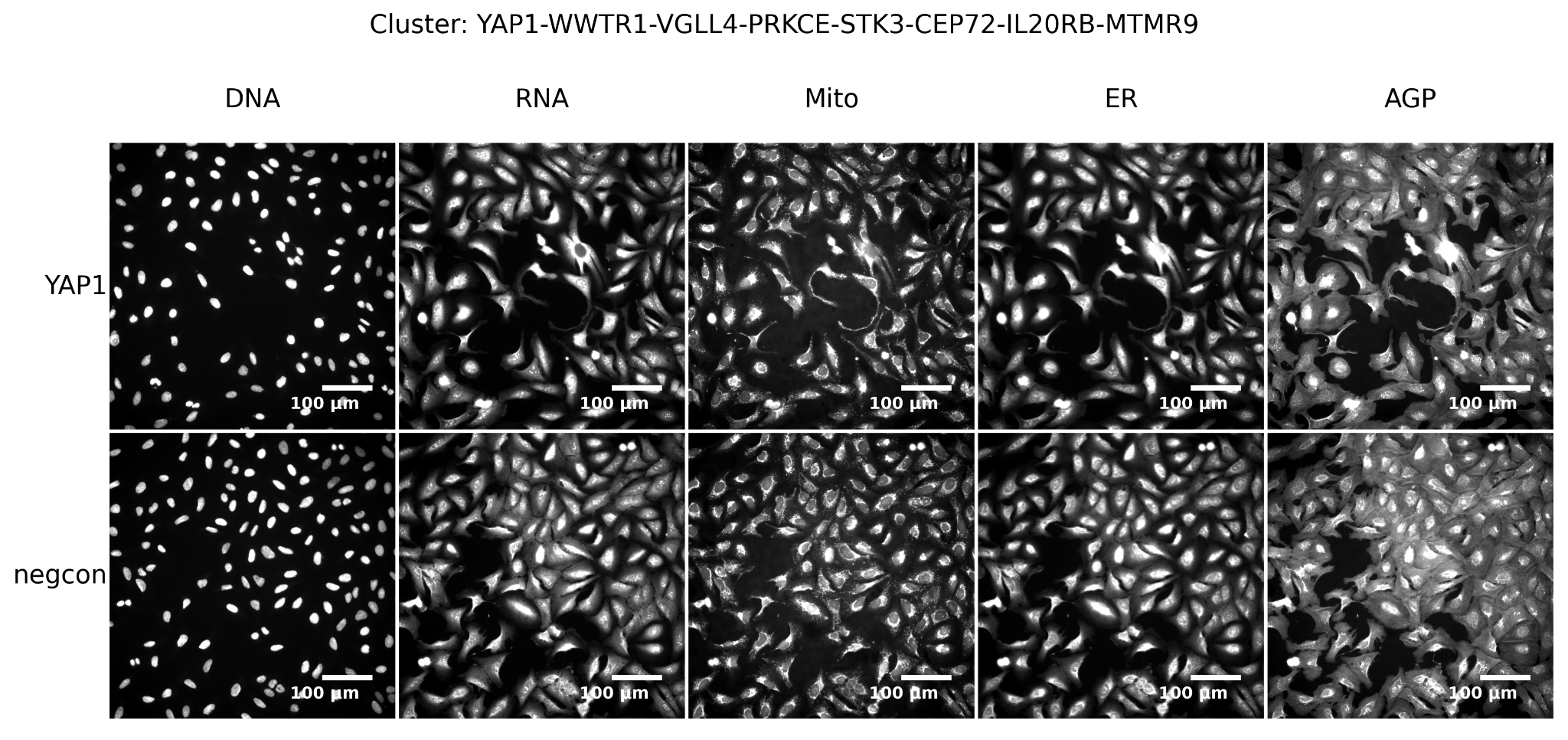

*Supplementary Figure 31**:* ***Example images from the YAP1 cluster in the ORF dataset.*** *Representative Cell Painting images for a randomly selected gene within the cluster (top row) and a randomly selected negative control (bottom row).*

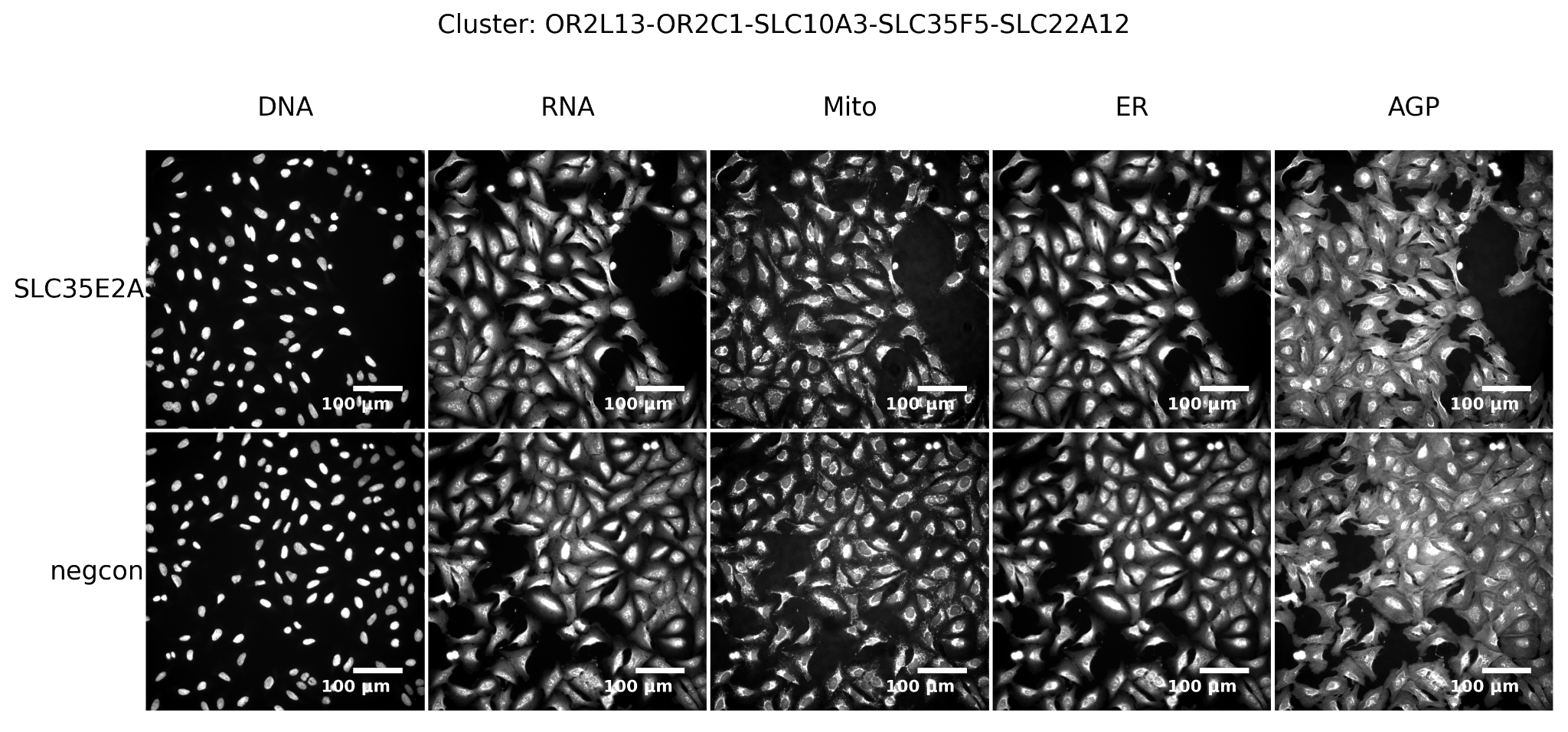

*Supplementary Figure 32**:* ***Example images from the SLC-OR cluster in the ORF dataset.*** *Representative Cell Painting images for a randomly selected gene within the cluster (top row) and a randomly selected negative control (bottom row).*

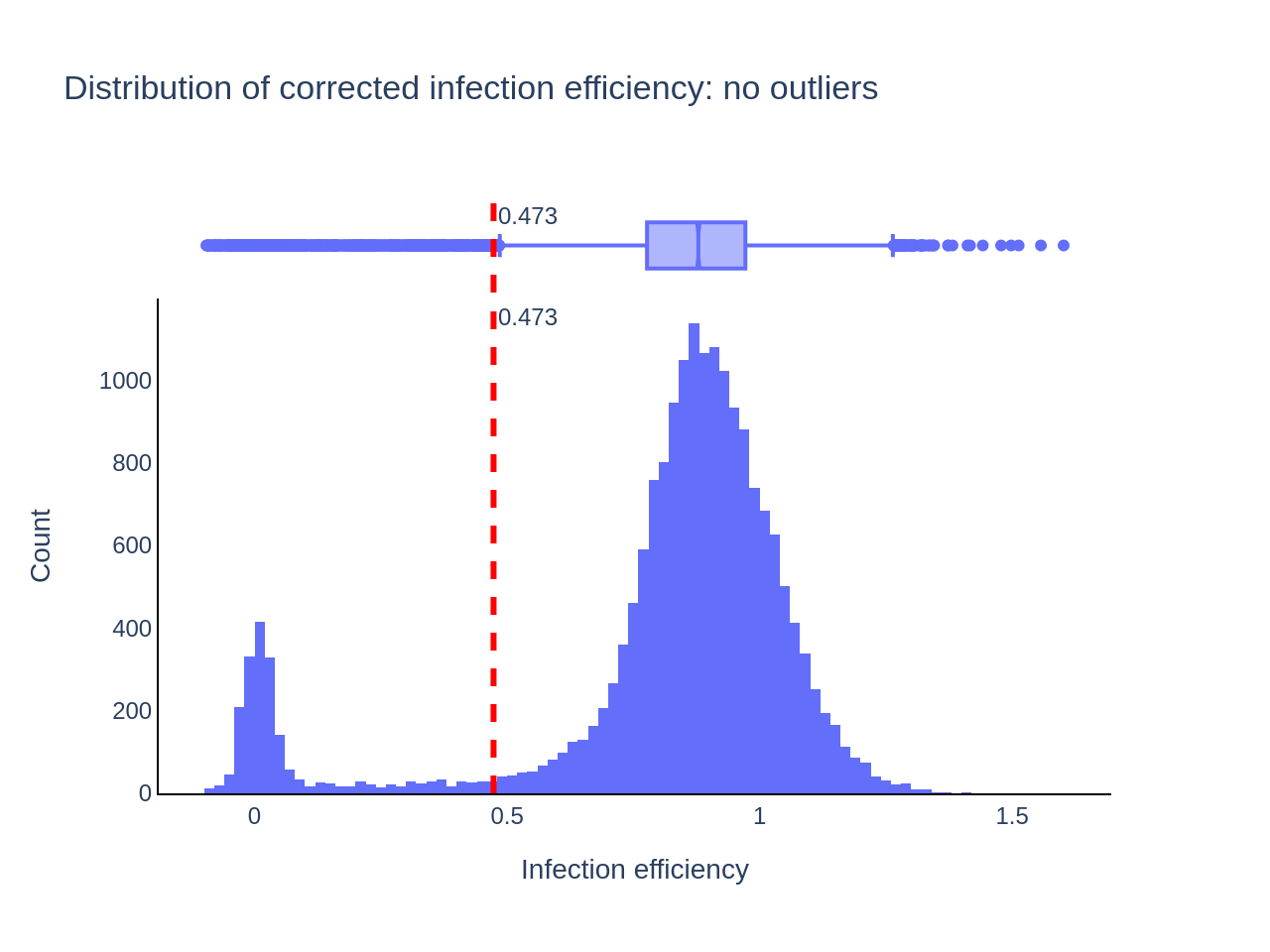

*Supplementary Figure 33**:* ***Histogram of infection efficiencies of ORF reagents.*** *Infection efficiency calculated using CellTiter-Glo® cell viability assay for all ORF reagents is plotted as a histogram. The threshold for low infection efficiency was determined by Otsu thresholding. Outliers were removed before plotting the histogram.*

| **Modality** | **Number of reagents distinguishable from negative control** | **Number of genes distinguishable from negative control** |
| --- | --- | --- |
| ORF | 7817 | 7031 |
| CRISPR | 5546 | 5546 |

*Supplementary Table 1**:* ***Phenotypic activity of ORF and CRISPR regents and genes.*** *The number of reagents and genes in the ORF and CRISPR data that have a phenotype is shown. Unlike the CRISPR reagents (which are pools of guides against each gene), there are sometimes multiple ORF reagents against many genes.*

| **Gene label** | **ORF** | **CRISPR** | **Common** |
| --- | --- | --- | --- |
| Disease association | 2 | 3 | 0 |
| CORUM complex | 103 | 369 | 54 |
| Wikipathway | 347 | 152 | 81 |
| HGNC Gene group | 196 | 89 | 13 |

*Supplementary Table 2**:* ***Phenotypic consistency.*** *Phenotypic consistency for different gene labels is shown. Some labels are retrieved in both the ORF and CRISPR datasets, while most retrieved labels are unique to the perturbation modality.*

| **Name** | **JCP2022_ID** | **InChIKey** |
| --- | --- | --- |
| AMG900 | JCP2022_037716 | IVUGFMLRJOCGAS-UHFFFAOYSA-N |
| LY2109761 | JCP2022_035095 | IHLVSLOZUHKNMQ-UHFFFAOYSA-N |
| Quinidine | JCP2022_050797 | LOUPRKONTZGTKE-UHFFFAOYSA-N |
| TC-S-7004 | JCP2022_012818 | CQKBSRPVZZLCJE-UHFFFAOYSA-N |
| NVS-PAK1-1 | JCP2022_064022 | OINGHOPGNMYCAB-UHFFFAOYSA-N |
| dexamethasone | JCP2022_025848 | GJFCONYVAUNLKB-UHFFFAOYSA-N |
| FK-866 | JCP2022_046054 | KPBNHDGDUADAGP-UHFFFAOYSA-N |
| aloxistatin | JCP2022_085227 | SRVFFFJZQVENJC-UHFFFAOYSA-N |

*Supplementary Table 3**:* ***Positive control compounds.*** *The first four are positive control compounds on the ORF plates, while the CRISPR plates contain all eight positive control compounds*

| *Method of machine learning model validation* | | | | | | | | | |
| --- | --- | --- | --- | --- | --- | --- | --- | --- | --- |
| Standard: Splitting the Gene–Function links in KG into training, validation and test sets | | | | | Time machine approach | | | | |
| **Model 1: Gene -> Gene Ontology Biological Process** | | | | | | | | | |
| *AUC* | *hits@10* | *hits@25* | *hits@50* |  | *AUC* | *AP* | *hits@10* | *hits@25* | *hits@50* |
| *0.99* | *0.543* | *0.689* | *0.787* |  | *0.82* | *0.83* | *0.118* | *0.196* | *0.275* |
| **Model 2: Gene -> Gene Ontology Molecular Function** | | | | | | | | | |
| *AUC* | *hits@10* | *hits@25* | *hits@50* |  | *AUC* | *AP* | *hits@10* | *hits@25* | *hits@50* |
| *0.98* | *0.8711* | *0.9258* | *0.9485* |  | *0.83* | *0.85* | *0.387* | *0.514* | *0.616* |
| ***Model 3: Gene -> Pathway*** | | | | | | | | | |
| *AUC* | *hits@10* | *hits@25* | *hits@50* |  | *AUC* | *AP* | *hits@10* | *hits@25* | *hits@50* |
| *0.99* | *0.820* | *0.911* | *0.954* |  | *N/A* | *N/A* | *N/A* | *N/A* | *N/A* |

*Supplementary Table* *4:* ***Validation of Graph Neural Network model for predicting gene function from the knowledge graph.*** *Time machine approach refers to using only those Gene Ontologies in the test set which appeared in the database after the knowledge graph creation date. We have not had this information for pathway databases.*
